# Ribosome remodeling drives translation adaptation during viral infection and cellular stress

**DOI:** 10.1101/2025.10.24.684008

**Authors:** Hsin-Yu Tsai, Luochen Liu, Rebecca H. Fleming, Julian Mintseris, Derrick Ekanayake, Xin Gu, Steven P. Gygi, Amy S.Y. Lee

## Abstract

The ribosome is the highly conserved molecular machine that decodes mRNAs during protein synthesis. While traditionally thought to consist of a uniform set of proteins, here we discover that ribosome composition is reprogrammed to adapt to intrinsic and external cellular perturbations. During infection by non-segmented negative-sense viruses, viral entry into cells recruits the large ribosomal subunit protein rpL40 to a noncanonical site on the small subunit of 80S ribosomes near the mRNA entry site. These specialized ribosomes preferentially bind viral mRNAs to drive enhanced viral protein synthesis that is critical for replication under host pressures. Unexpectedly, we find that viruses have co-opted this translation pathway from a previously unrecognized endogenous ribosome remodeling program in which metabolic stress alters ribosome structure to promote mRNA translation required for cell survival. Thus, ribosome remodeling is a conserved mechanism enabling dynamic protein synthesis across pathogen and cellular adaptation.

## Introduction

The ribosome is the core machinery responsible for decoding mRNA and catalyzing protein synthesis in all life^1^. Given its conserved and essential function, ribosomes were traditionally viewed as structurally and functionally invariant complexes. However, recent mass spectrometry suggests widespread differences in ribosome composition across tissues and developmental states^2-6^. Yet, there are few examples directly linking endogenous compositional changes to altered ribosome function, with most studies focused on protein loss or paralog switching. For example, release of rpS26 from *S. cerevisiae* ribosomes promotes translation of stress-response genes in response to osmotic or alkaline stress^7^, while tissue-specific mouse ribosomes containing the ribosomal protein paralog rpL39L-like are required for male fertility^4^. Beyond the limited documented changes to endogenous ribosome function, viruses can also modify host ribosomes through ribosome-targeting factors to preferentially translate viral mRNAs while suppressing antiviral protein synthesis^8,9^. For example, poxviruses phosphorylate the small ribosomal subunit protein RACK1 to direct selective translation of viral mRNAs^3^ and coronaviruses block host interferon responses by encoding a viral protein that sterically blocks the mRNA entry tunnel of 40S ribosome^10-12^. Overall, these observations raise key questions of how dynamic changes in ribosome composition may shape the proteome and enable adaptive responses in both cellular and viral contexts.

Here, we leverage viruses to discover a pathway that reprograms ribosome composition by incorporation of an additional copy of the core ribosomal protein rpL40 and drives translation adaptation critical during broad cellular perturbations. We identify membrane fusion during non-segmented negative-strand (NNS) RNA viral entry as a novel trigger for ribosome remodeling. The additional rpL40 is positioned at a non-canonical site on the small ribosomal subunit and enhances viral protein synthesis and fitness through directing ribosome binding to viral mRNA. We further find that this mechanism is paralleled during cellular serum starvation, revealing that both viruses and cells converge on ribosome remodeling to enable appropriate proteome adaptation and fitness. Altogether, these results illustrate an underexplored function of this essential machinery, beyond its core role in peptide bond synthesis, in selective mRNA translation. Furthermore, these findings highlight endogenous shifts in ribosome composition as a critical pathway for gene regulation, with broader implications on cell specification, adaptation, and pathogen infection.

## Results

### Ribosome remodeling is a conserved response to NNS virus infection

Viruses manipulate non-canonical host mechanisms to reprogram gene expression machinery, and defining these pathways has revealed previously unrecognized yet critical modes of cellular gene regulation^13-15^. However, whether they modify ribosome composition to reprogram the proteome to be advantageous for viral infection remains unclear^3^. To test this possibility, we leveraged NNS viruses, which include major human pathogens such as Ebola, rabies, and measles viruses. We infected human embryonic kidney (HEK) 293T cells with the prototype NNS virus vesicular stomatitis virus (VSV) and assessed ribosomal protein abundance on isolated 80S ribosomes by tandem mass tag-mass spectrometry (TMT-MS) (Fig. 1A, Table 1). As expected, most ribosomal proteins remained unchanged upon infection; however we surprisingly identified 11 ribosomal proteins as significantly increased on ribosomes, including the large subunit proteins rpL40, rpL9, rpL32, rpL13, and rpL37, and the small subunit proteins RACK1, rpS21, rpS6, rpS3, rpS11, rpS12 (Fig. 1B). Several of the identified RPs are surfaced exposed, such as rpL9, rpL32, rpL40, rpS3, RACK1 and rpS12 (Fig. S1A), indicating that external accessibility of these proteins could facilitate dynamic ribosome remodeling without requiring new ribosome assembly.

**Figure 1.**
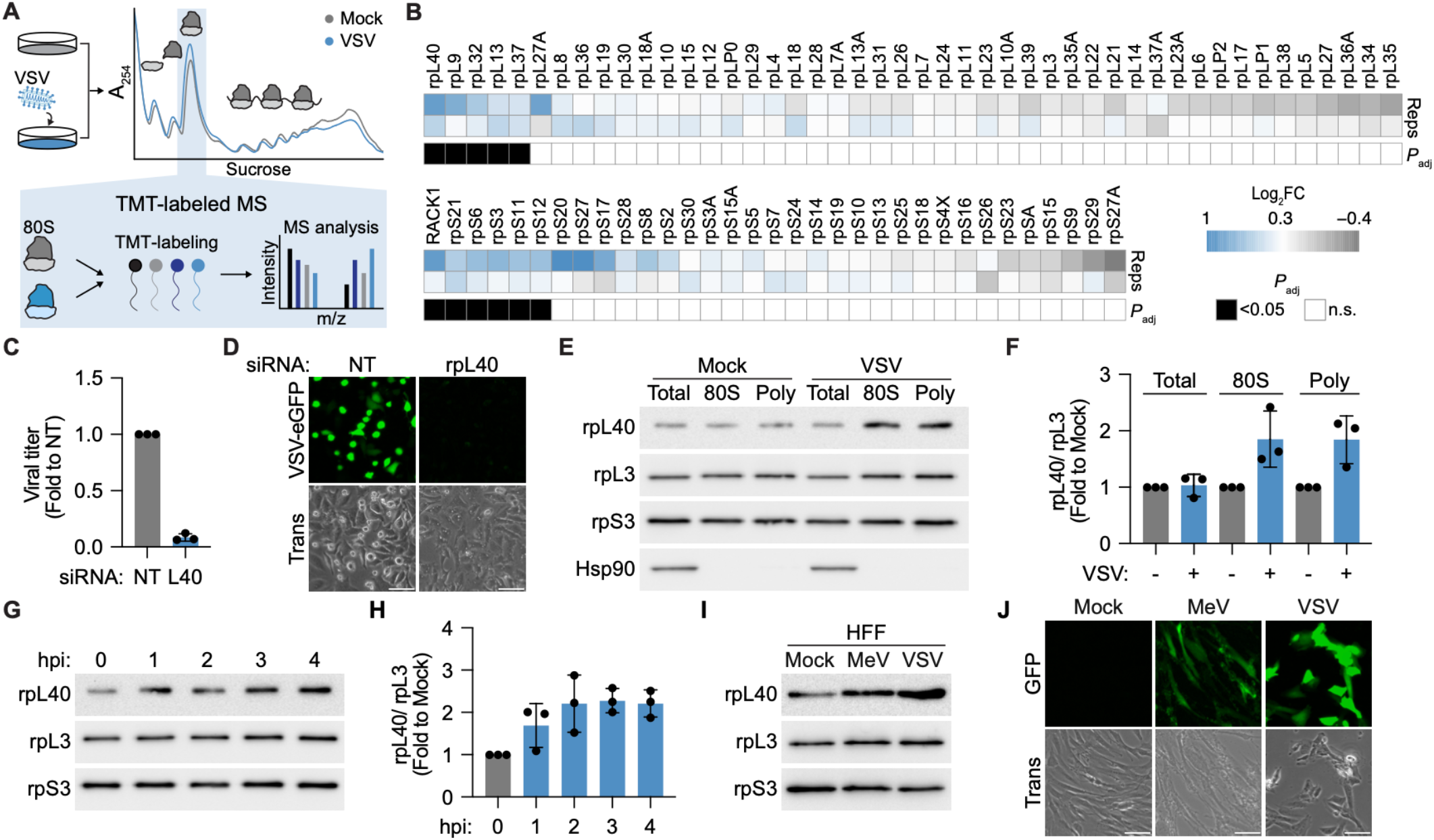
Ribosome remodeling is a conserved response to NNS virus infection. **A**. Schematic of analysis of ribosome heterogeneity during infection. HEK293T cells were mock or VSV-infected (MOI = 3, 3 hpi), 80S ribosomes were isolated from lysed cells by sucrose gradient ultracentrifugation, and ribosomal protein abundance was analyzed by TMT-MS. **B**. Heatmap showing relative changes in ribosomal protein (RP) abundance in 80S ribosomes from mock versus VSV-infected cells. RPs with statistically significant changes (Padj < 0.05) are marked with black squares, *n* = 2 biologically independent replicates, processed and measured in the same mass spectrometry acquisition. **C**. VSV titers in HeLa cells transfected with non-targeting (NT) or rpL40-targeting siRNA as measured by plaque assay (MOI = 10, 8 hpi). Results are normalized to NT-transfected cells and presented as mean fold change ± SD, *n* = 3 biologically independent samples. **D**. Fluorescence microscopy of HeLa cells transfected with NT or rpL40-targeting siRNA and infected with VSV-eGFP (MOI = 5, 5 hpi). Scale bar, 300 μm. Trans, transmitted light. **E**. Immunoblot and **(F)** quantification of rpL40 levels in total cytoplasmic lysates or 80S ribosome or polysomes isolated from mock or VSV-infected HEK293T cells (MOI = 3, 3 hpi). Hsp90 is a loading control for cytoplasmic protein. rpL3 and rpS3 are loading controls for purified ribosomes. **G**. Immunoblot and (**H**) quantification of rpL40 levels on purified 80S ribosomes during a VSV infection timecourse (MOI = 3). Results in (F)–(H) are represented as rpL40 levels normalized to rpL3 and presented as mean fold change ± SD relative to mock-infected cell samples, *n* = 3 biologically independent samples. **I**. Immunoblot of rpL40 levels on purified 80S ribosomes isolated from human foreskin fibroblasts (HFF) infected with measles virus (MeV, MOI = 10, 5 hpi) or VSV (MOI = 5, 5 hpi). **J**. Fluorescence microscopy of HFF cells infected with VSV-eGFP (MOI = 5, 24 hpi) or MeV-GFP (MOI = 10, 24 hpi). Scale bar, 300 μm. Results of (D), (E), (G), (I), and (J) are representative of *n* = 3 biologically independent samples.

We observed the greatest change in ribosome occupancy upon infection by the large ribosomal subunit protein rpL40. RpL40 is encoded as a ubiquitin–ribosomal protein fusion from the *UBA52* gene and is proteolytically cleaved off the ubiquitin to form the mature ribosomal protein L40 protein^16^. Notably, rpL40 was previously identified as a conserved host factor essential for replication of many NNS viruses, including measles virus, rabies virus, and Newcastle disease virus^17^ (Fig. 1C,D, Fig. S1B). In agreement with the mass spectrometry results, rpL40 levels increased 2-fold on 80S ribosomes and polysomes upon VSV infection as measured by immunoblotting (Fig. 1E,F). This shift in ribosome composition occurred rapidly, within the first hour after infection (Fig. 1G,H), and without an increase in total cytoplasmic rpL40 levels (Fig. 1E). Furthermore, ribosome remodeling upon VSV infection was conserved across multiple immortalized and primary cell types, including HeLa, A549, and human foreskin fibroblasts (HFF) (Fig. S1B). In addition, rpL40 increased on 80S ribosomes isolated from HFF cells infected with measles virus (Fig. 1I,J). Therefore, remodeling of rpL40 levels is a conserved ribosomal response to NNS virus infection across diverse cell types.

### Membrane fusion during virus entry induces rpL40 ribosomal occupancy

RpL40 occupancy on ribosomes increases rapidly upon VSV infection, suggesting that an early stage of infection such as viral entry or primary transcription induces ribosome remodeling (Fig. 2A). To determine the responsible step, we transfected purified VSV ribonucleoprotein (RNP) cores to initiate replication while bypassing viral entry. RpL40 occupancy was not induced by RNP transfection, revealing that viral entry is critical for ribosome remodeling (Fig. 2B,C, Fig. S2A). We next performed systematic perturbation to identify how viral entry triggers ribosome remodeling. VSV enters cells via clathrin-mediated endocytosis. Blocking internalization with Dynasore, which inhibits dynamin-mediated scission of clathrin-coated pits^18^, prevented increased rpL40 occupancy, indicating a post-internalization step is required for this process (Fig. 2D,E). As VSV has a large, bullet-shaped structure (70× 200 nm)^18^, approximately three times the size of a typical clathrin-coated vesicle, we tested if viral physical properties contribute to ribosome remodeling. Viral shape and size do not contribute to this change, as a spherical (∼80 nm) lentivirus pseudotyped with VSV-G also increased rpL40 occupancy (Fig. 2F).

**Figure 2.**
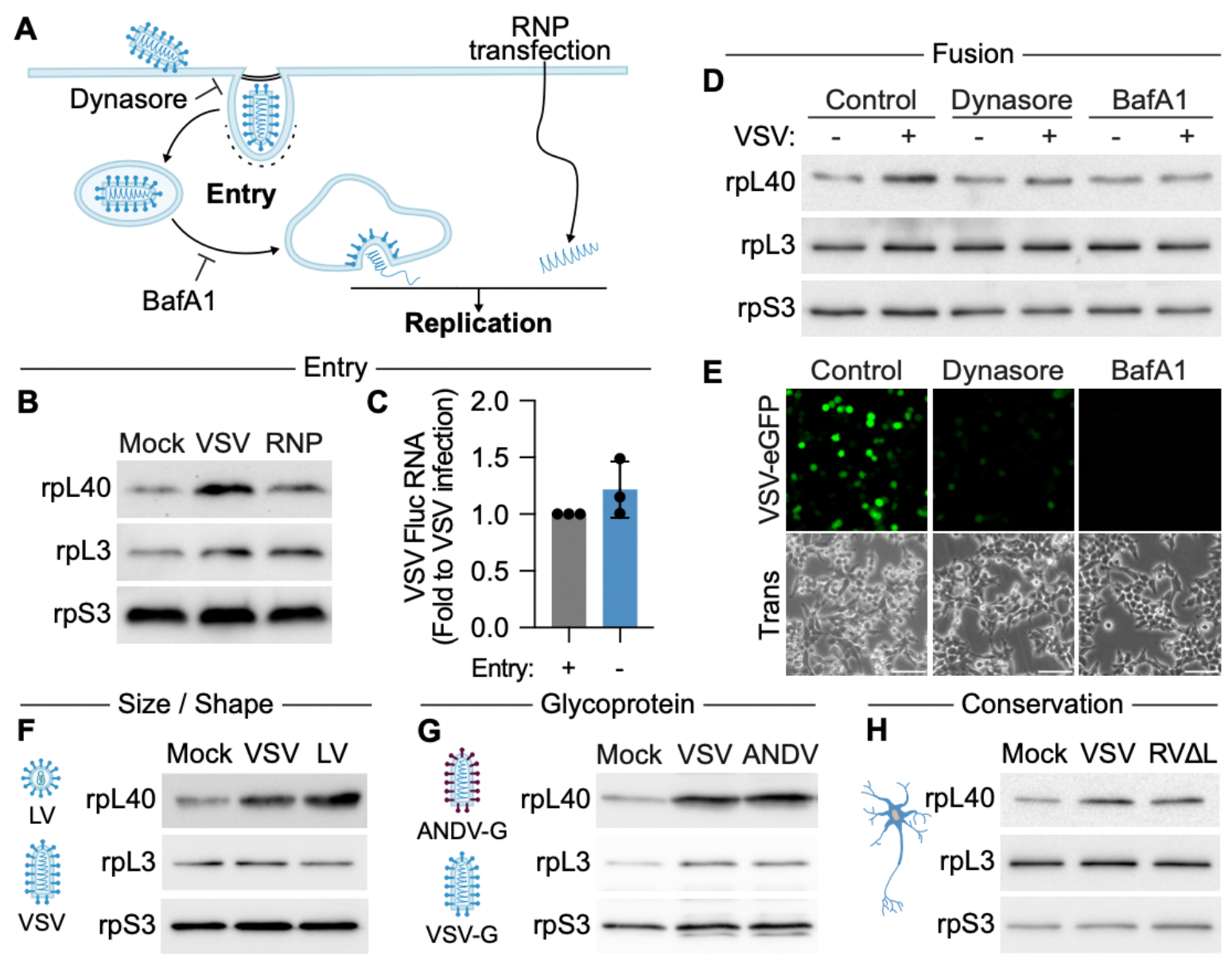
Membrane fusion during virus entry induces rpL40 ribosomal occupancy. **A**. Schematic of VSV entry and replication. **B**. Immunoblot of rpL40 levels on 80S ribosomes isolated from VSV-infected HEK293T cells or cells where viral entry is bypassed by RNP transfection. **C**. Quantification of VSV mRNA levels as measured by qRT-PCR and presented as mean ± SD normalized to VSV infection from *n* = 3 biologically independent samples. **D**. Immunoblot of rpL40 levels on 80S ribosomes isolated from mock or VSV-infected cells treated with inhibitors of viral internalization (Dynasore) or viral fusion (BafA1, Bafilomycin A1). **E**. Fluorescence microscopy of HEK293T cells infected with VSV-eGFP (MOI = 3, 4 hpi). Scale bar, 300 µm. Trans, transmitted light. **F**. Immunoblot of rpL40 levels on 80S ribosomes following infection with viruses of different size and morphology. LV, lentivirus pseudotyped with VSV-G. **G**. Immunoblot of rpL40 levels on 80S ribosomes following infection with VSV expressing endogenous or Andes virus (ANDV) glycoproteins. **H**. Immunoblot of rpL40 levels on 80S ribosomes isolated from primary mouse cortical neurons infected with VSV (MOI = 3, 3 hpi), or a polymerase-deficient Rabies virus (RVΔL, MOI = 10, 5 hpi). Result is representative of *n* = 2 biologically independent samples. Results of (B), (D), (E–H) are representative of *n* = 3 biologically independent samples.

Following endocytosis, virions undergo low pH-dependent fusion in early endosomes to allow for genome release into the cytosol. Blocking endosomal acidification during entry with the vacuolar proton pump inhibitor Bafilomycin A prevented ribosome remodeling, demonstrating that viral fusion is required for this process (Fig. 2D,E). Notably, the specific viral protein mediating fusion is not critical, as recombinant VSV expressing a highly divergent glycoprotein from the bunyavirus Andes virus (ANDV)^19,20^ also induced rpL40 levels on 80S ribosomes (Fig. 2G). To examine if the cellular compartment where viral fusion occurs contributes to ribosome remodeling, we leveraged the low pH-dependent fusion of the VSV glycoprotein. VSV particles were attached to cells at 4°C and then virus–cell fusion at the plasma membrane was activated by shifting the pH to 5.5 in the presence of Bafilomycin A to block endosomal fusion^18^. The specific localization of viral fusion is not critical for ribosome remodeling as fusion at the plasma membrane increased rpL40 occupancy (Fig. S2B,C). In agreement with a conserved induction mechanism of ribosome remodeling across NNS viruses, infection of primary cortical neurons with a polymerase-deletion strain of rabies virus (ΔL-RABV)^21^ that can enter cells via fusion but cannot replicate also increased rpL40 levels on 80S ribosomes (Fig. 2H, Fig. S2D,E). Collectively, these results reveal a novel regulatory mechanism at the host-virus interface, in which viruses remodel ribosomes prior to viral replication, thereby reprogramming the core protein synthesis machinery at the earliest stage of infection. Importantly, as membrane fusion is a conserved and fundamental step for the entry of all enveloped viruses, virus-specific signaling during fusion could be a mechanism to trigger distinct cellular priming of the ribosome and tune the host protein synthesis machinery for specific infection needs.

### Ribosome remodeling localizes rpL40 to a non-canonical site on the small ribosomal subunit

RpL40 is located at the solvent-exposed surface of the 60S subunit, near the GTPase activation center^22^. While our findings reveal that rpL40 levels on the 80S double upon viral infection, absolute mass spectrometry data indicate near-full stoichiometry of rpL40 in polysomes^5^, suggesting the additional copy of rpL40 could localize to a non-canonical position. To determine the rpL40 protein interface on remodeled ribosomes, we performed chemical crosslinking and mass spectrometry (XL-MS)^23^. Purified 80S ribosomes from VSV-infected cells were crosslinked using BS3, a homobifunctional chemical crosslinker that covalently links primary amines spatially proximal within ∼ 11.4 Å. (Fig. 3A, Table 2). Two LC-MS runs were searched with the PIXL search engine^24^, identifying 780 unique cross-linked peptides at 1% false discovery rate (671 unique cross-linked position pairs) across the whole ribosome. These included reproducible, high-confidence crosslinks involving rpL40, consisting of four peptides from three rpL40-crosslinked proteins (Fig. 3B). All crosslinks mapped to the α-amino group on the N terminal isoleucine residue of rpL40, consistent with existing structural data showing that the N terminal α-helix is surface-exposed while the central zinc-finger domain forms extensive contacts with the ribosomal RNA^22^. In agreement to the known location on the large ribosomal subunit, rpL40 crosslinked with rpL9 (K174, K184) and rpLP0 (K134), which are proximal to the canonical rpL40 location on large subunit by the P-stalk and GTPase activation center (Fig. 3C). Unexpectedly, we also identified crosslinks with the small ribosomal subunit protein rpS3 at positions K62 and K108, suggesting a novel interaction site of rpL40 on the small subunit at the mRNA entry tunnel (Fig. 3B,C, Fig. S3A). rpS3 is a conserved core protein of the 40S ribosomal subunit that mediates translation initiation by binding to mRNA and stabilizing the preinitiation complex^25^. K108 is positioned near the mRNA entry channel and involved in initiation fidelity, while K62 is critical for proper structural rearrangements of the 48S preinitiation complex required for the transition to elongation^25-27^. Using molecular structure prediction, we modeled an interaction pocket at the entry of the mRNA-binding tunnel that could feasibly position rpL40 by rpS3 as supported by the proximity information provided by XL-MS (Fig. S3A).

**Figure 3.**
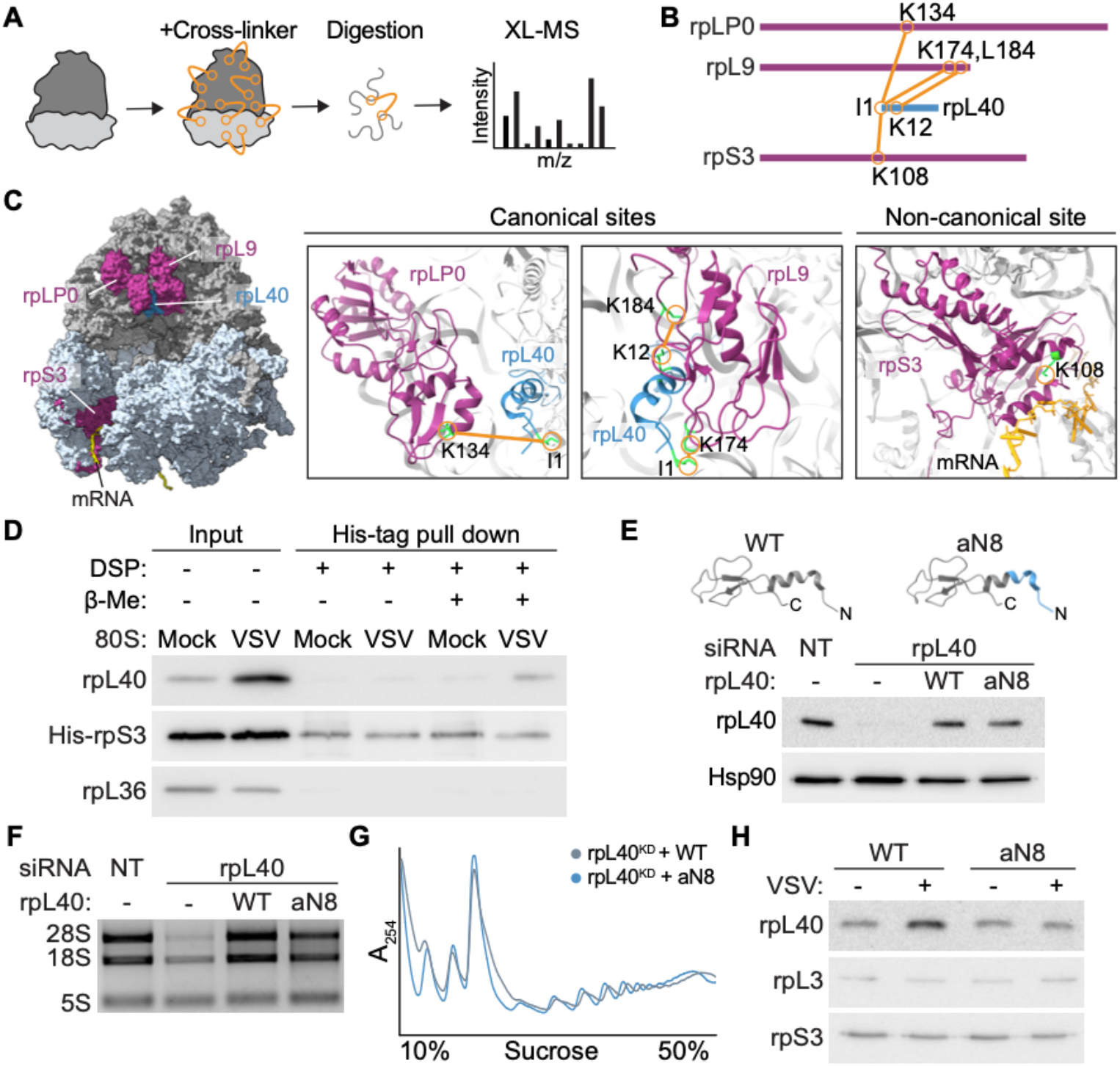
Ribosome remodeling localizes rpL40 to a non-canonical site on the small ribosomal subunit. **A**. Schematic of cross-linking mass spectrometry (XL-MS) workflow. **B**. Interaction network of rpL40 with ribosomal proteins on 80S ribosomes isolated from VSV-infected HEK293T cells as identified by XL-MS (*n* = 2 biologically independent samples). Proteins are represented by lines drawn to scale; each orange line represents one identified crosslink pair. **C**. Structural model of the 80S ribosome highlighting the positions of rpL40-interacting RPs and crosslinking sites. **D**. Crosslinking-immunoblot validation of the noncanonical interaction between rpL40 and rpS3 on 80S ribosomes isolated from mock or VSV-infected cells stably expressing His-tagged rpS3. DSP, crosslinker; β-Me, 2-mercaptoethanol. **E**. Immunoblot of cytoplasmic lysates from HeLa cells engineered to exclusively express wild type (WT) or N-terminal archaeal-chimeric (aN8) rpL40 by siRNA knockdown and stable gene expression rescue. **F**. Agarose gel electrophoresis of rRNA and (**G**) polysome profiles from cells following rpL40 knockdown and with rescue by WT or aN8 rpL40. **H**. Immunoblot of rpL40 levels on 80S ribosomes isolated from mock or VSV-infected cells exclusively expressing WT or aN8 rpL40. Results of (D–H) are representative of *n* = 3 biologically independent samples.

To biochemically validate this novel rpL40 binding site, we generated a HEK293T cell line stably expressing rpS3 with a C-terminal His-tag (His-rpS3) (Fig. 3D, Fig. S3B). We purified 80S ribosomes from mock or VSV-infected cells and crosslinked proteins using the thiol-cleavable amine-reactive crosslinker dithiobis (succinimidyl propionate) (DSP). As the ribosome is a tightly associated multi-protein complex, to isolate proteins directly interacting with rpS3, we applied high-stringency urea treatment to dissociate non-covalent interactions and used affinity purification to selectively capture His-tagged rpS3 and crosslinked partners. With intact DSP crosslinks, the rpS3–rpL40 interaction was not detectable, likely because simultaneous crosslinking of rpS3 to other proteins would form a large complex that cannot be resolved by electrophoresis. Upon treatment with 2-mercaptoethanol to release crosslinked proteins, we observed an interaction between rpL40 and rpS3 (Fig. 3D). Importantly, this interaction was specific as rpS3 did not crosslink to another large ribosomal subunit protein rpL36 (Fig. 3D), and was only captured in ribosomes isolated from VSV-infected cells, consistent with increased rpL40 occupancy during infection (Fig. 1E).

As the N-terminus of rpL40 crosslinks to rpS3, we hypothesized this region is critical for regulation of ribosome remodeling. We engineered a rpL40 mutant (aN8) where the N terminal 1-8 amino acids were replaced with the corresponding region from the archaeal *S. solfataricus* homolog^28^, therefore retaining the native α-helical structure while reducing amino acid identity to 25% to perturb interaction surfaces (Fig. 3E, Fig. S3C). In agreement that this mutant does not disrupt overall folding, HeLa cells exclusively expressing either wild-type or aN8 mutant rpL40 exhibited similar rRNA maturation, polysome profiles, and assembly of rpL40 into 80S ribosomes under homeostatic conditions (Fig. 3F–H, Fig.S3D). However, upon VSV infection, cells expressing the aN8 mutant were no longer able to increase rpL40 occupancy on ribosomes, indicating the rpL40 N-terminus is a key regulatory element in controlling ribosome remodeling (Fig. 3H). Thus, viral infection remodels ribosomes by directing rpL40 through the N-terminus to a non-canonical site near rpS3 at the mRNA entry tunnel.

### Increased rpL40 ribosomal occupancy promotes viral fitness under restrictive host conditions

NNS viral mRNAs, including those of VSV, structurally resemble host mRNAs by containing both a 5′ cap and a polyadenylated tail^29,30^. Despite this similarity, NNS viruses target host translation initiation factors to suppress the immune response^31^. Furthermore, for many NNS viruses, the 5′ UTRs of transcripts encoding replication proteins are too short to allow for conventional 40S scanning^32^. These features suggest NNS viral mRNA translation utilizes mechanisms distinct from canonical host mRNA translation. Given the proximity of rpL40 to the mRNA entry tunnel, we hypothesized ribosome remodeling facilitates ribosome engagement of viral mRNA during infection. Using purified 80S ribosomes from VSV-infected cells and radiolabeled VSV mRNAs *in vitro* transcribed by detergent-disrupted virions^33^, we found that 80S ribosomes directly bound viral transcripts (Fig. 4A,B). Importantly, this interaction was mediated by rpL40 as addition of rpL40-targeting antibodies, but not an isotype-matched IgG, blocked 80S–RNA binding (Fig. 4A,B). Ribosome binding was also specific to viral mRNAs, as addition of excess unlabeled VSV mRNA but not a cellular *ActB* mRNA competed away this interaction (Fig. 4C,D). Thus, remodeled ribosomes with higher rpL40 occupancy directly and selectively bind to VSV mRNAs.

**Figure 4.**
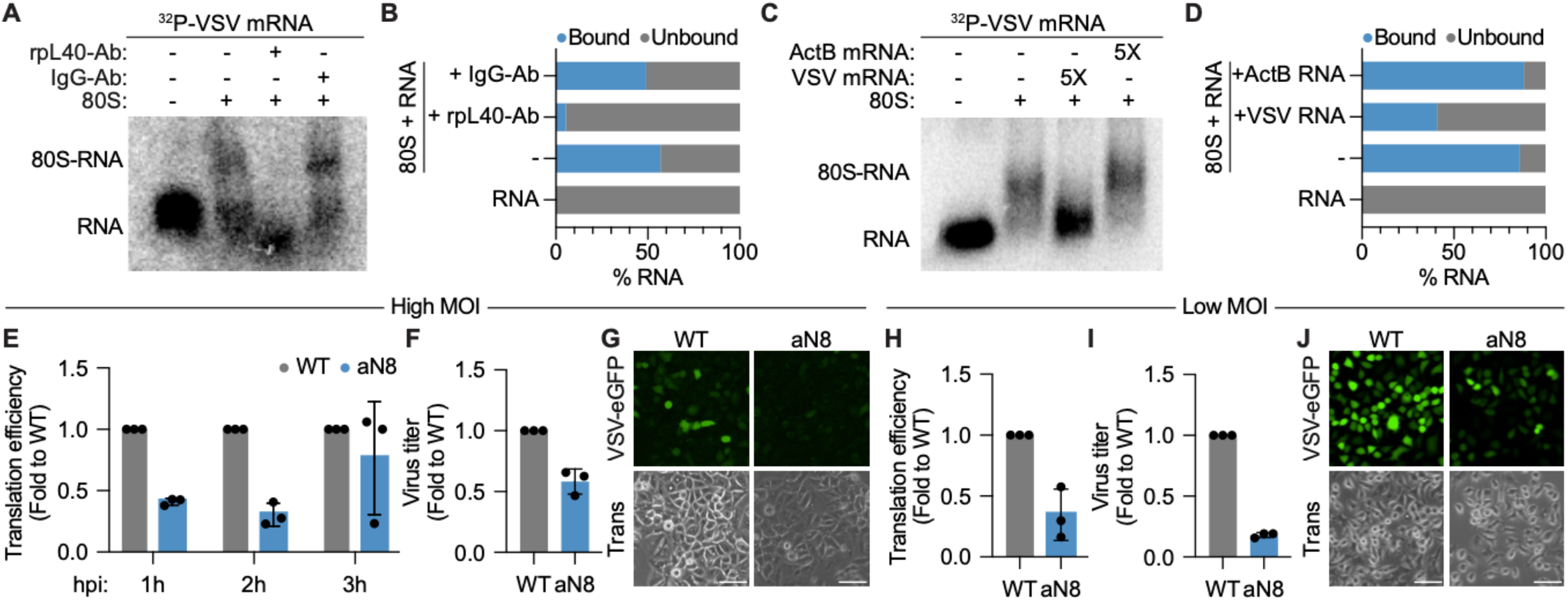
Increased rpL40 ribosomal occupancy promotes viral fitness under restrictive host conditions. **A**. Native gel shift assay and (**B**) quantification of ^32^P-labeled VSV mRNA binding to purified 80S ribosomes from VSV-infected cells. Addition of rpL40-targeting antibody, but not a control IgG, disrupts 80S–RNA complex formation. **C**. Native gel shift assay and (**D**) quantification of ^32^P-labeled VSV mRNA–80S complex formation in the presence of 5-fold molar excess of unlabeled VSV or cellular *ActB* RNA. **E**. VSV-*Fluc* mRNA translation efficiency during a high MOI infection timecourse in A549 cells exclusively expressing wild type (WT) or aN8 (occupancy mutant) rpL40 (MOI = 5). Translation efficiency was calculated as the ratio of protein levels normalized to *Fluc* mRNA levels, as measured by luciferase activity or qRT-PCR, respectively. **F**. VSV titers in A549 cells expressing WT or aN8 rpL40 as measured by plaque assay (MOI = 5, 5 hpi). **G**. Fluorescence microscopy of A549 cells expressing WT or aN8 rpL40 infected with VSV-eGFP. Scale bar, 300 µm. Trans, transmitted light. (MOI = 5, 4 hpi). **H**. VSV-*Fluc* mRNA translation efficiency, (**I**) VSV titers, and (**J**) fluorescence microscopy under low MOI viral spread conditions in A549 cells exclusively expressing wild type (WT) or aN8 (occupancy mutant) rpL40 (MOI = 0.01, 16 h). Scale bar, 300 µm. The results in (E) and (H) are presented as mean fold change ± SD in translation efficiency relative to WT cells, *n* = 3 biologically independent samples. The results in (F) and (I) are normalized to WT cells and presented as mean fold change ± SD, *n* = 3 biologically independent samples. The results in (B) and (C) are mean from = 3 biologically independent samples. The results of (A), (C), (G), and (J) are representative of *n* = 3 biologically independent samples.

To investigate the functional relevance of ribosome remodeling on viral translation, we next compared translation efficiency in HeLa cells expressing wild type or the aN8 rpL40 occupancy mutant by leveraging a recombinant VSV encoding firefly luciferase that allows for matched quantification of protein and RNA levels. Unexpectedly, we did not observe any difference in viral mRNA translation efficiency between the two cell lines (Fig. S4A). Viruses often evolve adaptations that are only advantageous under specific cellular contexts, such as immune pressure, and may have no detectable effect under permissive conditions^34,35^. As HeLa cells possess a highly attenuated antiviral response^36^, we hypothesized ribosome remodeling might contribute to context-dependent viral fitness. In support, A549 lung epithelial cells, which possess a functional innate immune system^37^, exhibited decreased translation efficiency at earlier infection timepoints when ribosome remodeling was blocked (Fig. 4E–G, Fig. S4B). Notably, viral mRNAs compose less than 1% of the total transcript pool early in infection, providing a challenge for viral mRNAs to compete for host translation machinery^38^. This defect in translation efficiency and viral output was further exacerbated under low MOI infection conditions, where viral spread is more constrained due to cytokine secretion from infected cells priming neighboring cells to create an environment that impairs efficient early gene expression^39,40^ (Fig. 4H–J). Therefore, virus-induced ribosome remodeling to increase rpL40 occupancy enhances viral translation during early infection and under restrictive host conditions.

### Ribosome remodeling is required for survival during cellular starvation

Viruses frequently exploit host cell stress response mechanisms to reprogram the translational landscape during infection, including core pathways such as the integrated stress response, DNA damage, mTOR signaling, and oxidative stress responses^41-44^. In yeast, rpL40 is required for cell survival in response to inappropriate mitochondrial protein import and was previously shown to regulate DNA-damage response mRNAs^17,45,46^. Given its potential role in stress adaptation, we hypothesized that remodeling of rpL40 on ribosomes is a conserved strategy used by both viruses and cells. Notably, 6 h of serum starvation led to increased rpL40 occupancy in HeLa cells, and this was abolished in cells expressing the aN8 mutant, indicating the same mode of ribosome remodeling is conserved between viral infection and nutrient stress (Fig. 5A,B). To determine how ribosome remodeling contributes to cellular gene regulation during stress, we sequenced total RNA and highly translating, polysome-associated mRNAs in wild type and aN8 rpL40-expressing cells. Supporting a role of ribosome remodeling in specialized translation regulation, RNA-seq revealed comparable transcript abundances between the two cell lines under both complete media and serum starvation conditions (Fig. S5A). mRNA translation was largely unaffected in complete media, in agreement with the absence of ribosome remodeling under homeostasis (Fig. 5C). In contrast, upon serum starvation, we identified 155 mRNAs that showed more than a two-fold reduction in translation efficiency in cells unable to increase rpL40 occupancy on ribosomes (Fig. 5C, Table 3). Gene ontology analysis revealed these mRNAs were enriched in essential metabolic and intracellular transport pathways (Fig. 5D), and included mRNAs encoding proteins critical for cellular adaptation to stress, such as *ESRRA* (Estrogen related receptor, alpha)^47^ and *TFEB* (Transcription Factor EB)^48^, which are transcription factors involved in survival during nutrient deprivation; and *SNX17*, a sorting nexin involved in recycling autophagosomal components^49^ (Fig. 5E, Fig. S5B, Table 3). Consistent with a critical role of increased rpL40 occupancy for selective translation during serum starvation, *SNX17* mRNA could not efficiently associate with highly translating polysome fractions in cells expressing aN8 rpL40, despite unchanged translation of the control *PSMB6* mRNA (Fig. 5E,F, Fig. S5B).

**Figure 5.**
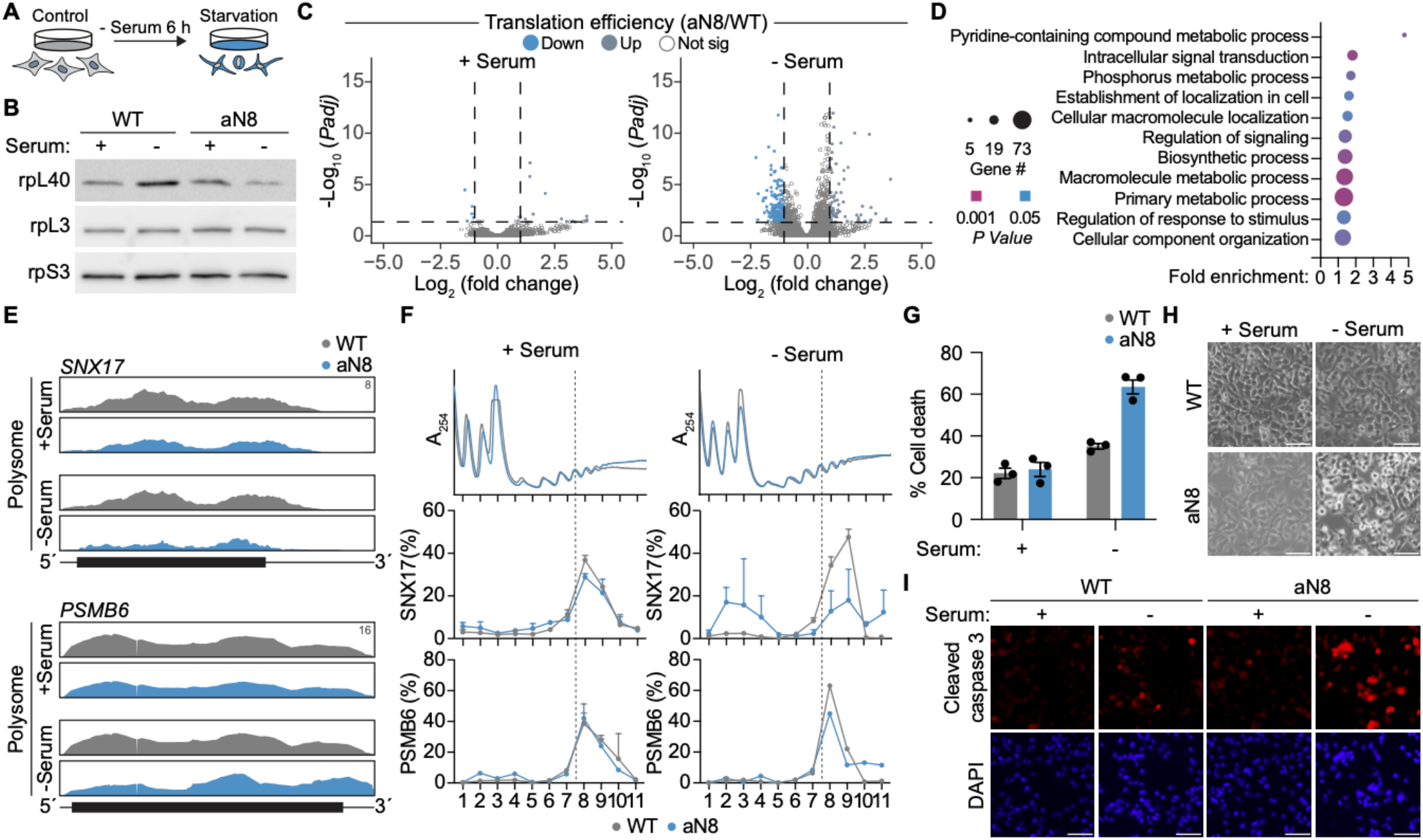
Ribosome remodeling is required for survival during cellular starvation. **A**. Schematic of serum starvation experiments. Serum starvation was performed by growing cells in serum-free media for 6 h. **B**. Immunoblot of rpL40 levels on 80S ribosomes isolated from WT or aN8 rpL40 HeLa cell lines grown under complete media or serum-starved conditions for 6 h. **C**. Volcano plot comparing the fold change in translation efficiency with adjusted *P* value of mRNAs in HeLa cells expressing WT versus aN8 rpL40 under control or serum starvation conditions. **D**. Gene ontology analysis of mRNAs translationally downregulated in serum-starved cells expressing aN8 rpL40 compared to WT rpL40. **E**. Read mapping to *SNX17* and *PSMB6* from Polysome-Seq under control or serum starvation conditions. The annotated y-axis maximum is equivalent for all samples. Transcript architecture is represented as follows: UTRs (thin line), coding regions (thick line). **F**. Association of *SNX17* and *PSMB6* mRNA with translating ribosomes in rpL40 WT or mutant cell lines under control or serum starvation conditions. mRNA abundance is expressed as a percentage of total transcript levels and plotted as mean ± SEM from two technical qRT-PCR replicates. Results are representative *n* = 2 biologically independent samples. **G**. Analysis of cell death in rpL40 WT or mutant cell lines. The results in (F) presented as mean ± SEM of *n* = 3 biologically independent samples. **H**. Images of cell morphology in rpL40 WT or mutant cell lines under control or serum starvation conditions. Scale bar, 300 μm. **I**. Immunofluorescence microscopy of cleaved-caspase 3 levels in rpL40 WT or mutant cell lines under control or serum starvation conditions. Nuclei are stained with DAPI. Scale bar, 300 μm. The results of (B), (H), (I) are representative of *n* = 3 biologically independent samples. Results of (C–E) are from *n* = 2 biologically independent samples.

Notably, we also identified key mRNAs involved in autophagosome-lysosome fusion and endosome recycling that required ribosome remodeling for efficient translation, including *HGS* (Hepatocyte Growth Factor-Regulated Tyrosine Kinase Substrate), a component of the ESCRT-I complex involved in endosomal sorting^50^; *VPS25* (Vacuolar Protein Sorting 25), part of the ESCRT-II complex that drives autophagosome maturation^51^; and *SPHK2* (sphingosine kinase-2), which generates sphingosine-1-phosphate to support lysosomal autophagic flux^52^. Cells which failed to induce rpL40 occupancy exhibited elevated levels of the autophagy marker LC3B-II, indicating that loss of ribosome remodeling-mediated translational control was sufficient to impair autophagy flux or autophagosome accumulation (Fig. S5C). In agreement with appropriate autophagy being critical for cells to adapt to nutrient starvation^53^, we additionally observed enhanced apoptotic cell death under serum starvation in aN8 mutant cells (Fig. 5G–I, Fig. S5C,D). Thus, cellular remodeling of ribosomes to increase rpL40 occupancy is essential for translation adaptation and survival during serum starvation, and NNS viruses have hijacked this endogenous stress response pathway to promote viral protein synthesis under host pressure.

## Discussion

The concept of ribosomal heterogeneity challenges the traditional view of ribosomes as passive machines in translation elongation^54-56^. However, the functional consequences of variations in ribosome composition remain poorly understood, thus limiting insights into if such dynamic changes to the core protein synthesis machinery are biologically significant. Here, our study reveals that cells increase rpL40 levels on ribosomes and remodeled ribosomes mediate selective mRNA translation. We identify a novel mechanism in which viral infection induces ribosome remodeling via fusion, and reveal a non-canonical mode of ribosome heterogeneity, where an additional copy of rpL40 is localized at an unanticipated site on the small ribosomal subunit. As both viral infection and serum starvation trigger the same ribosome remodeling, these findings reveal that viruses exploit an endogenous translation control mechanism for selective protein synthesis^15^.

RpL40 is considered a constitutive member of the large ribosomal subunit^57,58^. Our results reveal that during viral infection, an additional copy of rpl40 localizes proximal to rpS3 by the mRNA entry tunnel in the 40S ribosomal subunit, and that these ribosomes can bind directly to viral mRNAs. RpS3 plays a key role in control of translation initiation by stabilizing mRNA binding and ensuring accurate positioning of initiation factors for start codon^25,26,59^. Structural modeling of rpL40 and rpS3 predicts the C-terminus of rpL40 faces the mRNA entry tunnel, a location where this positively-charged lysine-rich terminus could enable interactions with specific transcripts at the RNA entry channel. Intriguingly, during SARS-CoV-2 infection, the viral protein Nsp1 interacts with rpS3 to block the mRNA entry channel and suppress host antiviral protein synthesis while selectively promoting viral mRNA translation^12,60,61^. Therefore, it is notable that viruses have independently evolved mechanisms to achieve transcript-specific translational control by targeting the same ribosomal region, and underscores the ribosome as a critical evolutionary interface in virus-host interactions^3,62,63^.

Our results show that ribosome remodeling occurs rapidly in response to cellular perturbations, including fusion during viral infection and intrinsic nutrient stress. What cellular signaling pathways could drive increased rpL40 occupancy? Notably, membrane fusion during infection requires actin remodeling to facilitate membrane bending and fusion pore formation^64^. Serum starvation also leads to altered cytoskeletal dynamics, especially through essential roles in autophagy such as phagophore formation and fusion with lysosomes^65,66^. Intriguingly, mass spectrometry analyses suggest that rpL40 transitions from high to low occupancy on ribosomes during mouse testis development, coinciding with the initiation of meiosis, a process dependent on dynamic actin remodeling^4,67^. These shared features suggest that actin remodeling could be a possible upstream signal that links membrane dynamics to ribosome remodeling. Understanding what are the signaling pathways that trigger remodeling, and if they are conserved across different cellular perturbations to coordinate shared translation adaptation, is an important focus of future research.

Here, we focused on rpL40 as a critical ribosomal protein required for viral fitness and cellular stress adaptation^17^. While genetic modulation of rpL40 occupancy is sufficient to have consequential impact on ribosome function, our mass spectrometry of 80S ribosomes during infection identified additional ribosomal proteins exhibiting dynamic occupancy changes upon infection. Notably, most of these proteins are located on the ribosome surface, consistent with greater accessibility to regulatory signals or remodeling without altering the core ribosomal functions of peptide bond formation. Indeed, evolutionary analyses show that while the ribosome core remains highly conserved, peripheral regions exhibit greater structural divergence^1^. Thus, the evolutionary plasticity of these surface components may contribute to their functional versatility in enabling ribosome remodeling and dynamic translational control in response to cellular conditions.

## Acknowledgements

We thank J. Guo, H. Kim, J. Vranken, M. Fels, R. Han, K. Erlenbeck, D. Hwang, S. Shao, C. Allard, N. Bellono, K. Chat, and members of the Lee lab for experimental advice, editing, or discussions. TMT-MS consultation and services were performed by the Thermo Fisher Center for Multiplexed Proteomics at Harvard Medical School. RNA-sequencing was performed by the Molecular Biology Core Facilities at Dana-Farber Cancer Institute, utilizing an Illumina NovaSeq X Plus funded by a National Institutes of Health SIG grant (1S10OD036228-01). This work was funded by grants to A.S.Y.L. from the Pew Biomedical Scholars Program, The G. Harold and Leila Y. Mathers Charitable Foundation, and the National Institutes of Health (1R35GM142527); grants to X.G. from the Aramont Fund for Emerging Science Research Award and a Damon Runyon Cancer Research Foundation Dale F. Frey Breakthrough Scientist Award (DFS 68-25); a grant to S.P.G. from the National Institutes of Health (GM067945); and R.H.F was funded by a Genetics Training Grant from the National Institutes of Health (GM007122).

## Author contributions

Conceptualization, HT, LL, RHF, ASYL; Investigation, HT, LL, RHF, JM, DE, XG; Data Curation, HT, ASYL; Writing – Original Draft, HT, ASYL; Writing – Review & Editing, HT, JM, LL, RHF, XG, DE, ASYL; Visualization, HT, JM, DE; Supervision, ASYL; Funding Acquisition, XG, SPG, ASYL

## Materials and Methods

### Resource Availability

Further information and requests for resources and reagents should be directed to and will be fulfilled by the lead contact, Amy S.Y. Lee.

### Materials Availability

Reagents and materials produced in this study are available from the lead contact pending a completed Materials Transfer Agreement.

### Cells and viruses

HEK293T, A549, HeLa, BHK21, Vero-6, and human foreskin fibroblast (HFF) cells were cultured in complete media (DMEM supplemented with 10% FBS (Biowest)). For serum starvation experiments, the media was removed and replaced with DMEM for 6 h. Mouse embryonic cortical neurons were prepared as previously described^70^. Briefly, cortices from wild-type C57BL/6 mouse embryos (Charles River Laboratories; 5–9 embryos, both sexes) at E16.5–E17 were dissected and enzymatically dissociated with papain (Sigma Aldrich, 10108014001). Digestion was stopped using ovomucoid (Worthington trypsin inhibitor), and the cells were gently triturated with a P1000 pipette, filtered through a 40-µm mesh, and plated onto poly-D-lysine (20 µg mL^-1^) and laminin (4 µg mL^-1^)-coated culture plates. Neurons were maintained in Neurobasal medium (Gibco) supplemented with 2% B27, 50 U mL^-1^ penicillin, 50 U mL^-1^ streptomycin, and 1 mM GlutaMAX, at 37°C in 5% CO_2_. At 3 days *in vitro*, neuronal culture was infected with VSV (MOI = 3, 3h) or ΔL-RABV (MOI = 10, 5h) and then harvested for polysome profiling and immunoblotting analyses. All animal procedures complied with protocols approved by the Harvard University Standing Committee on Animal Care and adhered to federal regulations.

Initial virus stocks were kind gifts from: rVSV-eGFP and rVSV-Fluc from S. Whelan (Washington University in St. Louis, St. Louis, MO); Measles-GFP virus^71^ from P. Duprex, University of Pittsburgh, Pittsburgh, PA); ΔL Rabies virus^21^ from I. Wickersham (Massachusetts Institute of Technology, Cambridge, MA); rVSV-ANDVg^19,20^ from K. Chandran (Albert Einstein College of Medicine, Bronx, NY). Vesicular stomatitis virus (VSV) expressing eGFP (rVSV-eGFP)^72^ or Fluc (rVSV-Fluc)^18^ or the Andes virus glycoprotein (rVSV-ANDVg)^19,20^ were amplified in BHK-21 cells as previously described^73^. Virus titer was determined by plaque assay on Vero E6 cells. Briefly, virus-containing media was serially diluted, and 200 µL of each dilution was used to infect Vero E6 cells at ∼90% confluency, seeded the day before in 6-well plates. Infection was carried out at 37 °C for 1 h with gentle rocking every 15 min. After adsorption, the media was removed, and 3 mL of sterile agar overlay (5% v/v FBS, 1× penicillin-streptomycin (Gibco), 0.292 mg mL^-1^ glutamine, 1× MEM with Earle’s salts, 0.12% w/v NaHCO_3_, 25 mM HEPES–KOH pH 7.5, and 0.25% w/v Oxoid agarose) was applied to each well. The overlay was solidified at room temperature for 15 min, and cells were incubated for ∼20 h at 37 °C. Plaques were fixed with 10% v/v formaldehyde in PBS for 15 min and stained with 0.05% w/v crystal violet in 10% v/v EtOH for 10 min. Lentivirus pseudotyped with VSV-G was rescued as previously described^74^. For viral entry studies, HEK293T cells were seeded in 10 cm plates to be at ∼70% confluency at the time of infection. To inhibit viral entry, HEK293T cells were pre-treated with 80 µM Dynasore inhibitor (HY-15304, MedChemExpress) or 100 nM Bafilomycin A1 (HY-100558, MedChemExpress) for 30 min before virus addition.

### VSV RNP transfection

VSV-Fluc viruses were concentrated from virus-containing media by centrifugation at 25,000 rpm for 3 h using a TH-641 rotor. The virus pellet was resuspended in NTE buffer (10 mM Tris–HCl pH 7.4, 100 mM NaCl, 1 mM EDTA). VSV RNPs were extracted as previously described^17^. Briefly, 600 µL purified and concentrated VSV-eGFP was treated with 300 µL 3× membrane-solubilization buffer (37.5 mM Tris–HCl pH 7.5, 15% v/v glycerol, 15 mM EDTA–KOH pH 8, 10.5 mM DTT, 0.3% Triton X-100, 1.5 M CsCl) on ice for 1 h, then loaded onto a 10–40% v/v glycerol gradient made in NTE buffer. The gradients were centrifuged using a TH-641 rotor at 38,000 rpm for 7 h at 4 °C. The RNP pellet was resuspended in NTE buffer and stored at –80 °C. For transfection, RNP was mixed with Lipofectamine 2000 at a 1 µg: 2.5 µL ratio in OptiMEM and incubated at RT for 15 min before being added to wells. Transfection efficiency was monitored by RT-qPCR using primers targeting the VSV N gene, compared to a VSV infection control.

### VSV fusion-infection

Low pH-induced fusion of VSV at the plasma membrane was performed as previously described^75^. HEK293T cells were seeded in 6-well plates coated with 0.1 mg mL^-1^ poly-D-lysine to be ∼70% confluent at time of infection. To attach VSV to the cell membrane, virus (MOI = 3) was added to the wells and the plate was centrifugated at 850 × g for 1 h at 4 °C. Control cells were centrifuged under the same conditions without virus. Cells were washed twice with cold PBS on ice and incubated with DMEM containing 10% FBS at pH 5.5 for 1 min at 37 °C before returning to ice. Media was replaced with standard complete DMEM containing 100 nM Bafilomycin A1 to block endosomal entry, and cells were incubated at 37 °C for 3 h before harvesting for ribosome analysis.

### Stable cell lines

To make the cell lines exclusively expressing rpL40 mutants, stable cell lines expressing WT or aN8 rpL40 were first created by lentivirus infection. For lentivirus production, HEK293T cells were seeded into 10 cm plates coated with 0.1 mg mL^-1^ poly-D-lysine a day before transfection, reaching ∼80% confluency at the time of transfection. A total of 9 µg PSPAX2 (packaging vector), 0.9 µg pMD2.G (VSV-G vector), and 9 µg of target gene constructs (described above) were mixed with 1 mL DMEM and PEI (PEI volume: DNA = 3:1), and incubated at RT for 15 min before being added to cells. After 24 h, media was replaced with DMEM containing 30% FBS and 25 mM HEPES-KOH pH 7.5. Media containing lentivirus was collected at 48 h post-transfection. Cell debris was removed by centrifugation at 500 × g for 5 min at RT. Virus-containing media was aliquoted and stored at –80 °C. For cell line establishment, cells were incubated with lentivirus and 1× polybrene for 24 h. After 24 h infection, media was replaced with complete media containing appropriate antibiotic(s), e.g., puromycin (1 µg mL^-1^) and/or hygromycin B (150 µg mL^-1^). HEK293T cells expressing His-tagged rpS3 were made by lentiviral infection as described above.

To knockdown endogenous rpL40 and allow for exclusive expression of the exogenous rpL40 ORFs in the stable cell lines, small interfering RNAs (siRNAs) were purchased from Dharmacon: siGENOME Non-Targeting siRNA #3 (D-001210-03-20) and siGENOME Human UBA52 siRNA (D-011794-02-0050). The lyophilized siRNA was resuspended in siRNA buffer (20 mM KCl, 6 mM HEPES–KOH pH 7.5, 0.2 mM MgCl_2_) and folded by heating at 95 °C for 3 min followed by slow cooling to 4 °C. siRNA was transfected into HeLa or A549 cells with a final working concentration of 75 nM using Lipofectamine 2000 (Invitrogen) as previously described^17^. Cells were incubated for 48 h to allow for gene knockdown before performing additional experiments.

### Cloning

Lentivirus transfer plasmids containing WT or aN8 rpL40 were constructed by ligating synthetic gene fragments into the N144-Hygro vector digested with BamHI and MluI. The lentivirus transfer plasmid expressing His-rpS3 was designed with a C-terminal 6X His-tag and GSGS linker. The construct was generated by amplification of rpS3 from HEK293T cDNA and cloned N144 as described above. The lentivirus transfer plasmid to rescue lentivirus pseudotyped with VSV-G for entry experiments was constructed by cloning GFP into the N144-Puro vector digested with BamHI and NotI. Primers used for cloning are listed in Table S1.

### Reverse transcription and quantitative PCR

RNA was isolated using Trizol Reagent (Invitrogen) following the manufacturer’s protocol. cDNA was synthesized from RNA by reverse transcription using random hexamers and MMLV M5 reverse transcriptase under standard conditions as previously described^74^. Quantitative PCR was performed using Luna Universal qPCR master mix (NEB) and primers listed in Table S1.

### Viral translation efficiency

Cells were infected with VSV-Fluc or transfected with VSV-Fluc RNP as indicated, and then collected and pelleted by centrifugation at 300 × g for 3 min at 4 °C. Cells were lysed with 1× Luciferase Cell Culture Lysis buffer (E153A, Promega) on ice for 5 min. For protein synthesis levels, relative luminescence units were measured using a Luciferase System kit (Genecopoeia) according to the manufacturer’s protocol and a Glomax Multi+ plate reader (Promega). For *Fluc* mRNA levels, total RNA extraction, cDNA synthesis, and quantitative PCR were performed as described above. Translation efficiency was determined by first quantifying *Fluc* mRNA levels using qPCR and normalizing to *GAPDH* mRNA levels. The luciferase activity was subsequently divided by the corresponding normalized mRNA quantity to calculate translation efficiency.

### Immunoblotting

Protein samples were mixed with 4× SDS loading buffer (200 mM Tris–HCl pH 6.8, 400 mM β-mercaptoethanol, 8% w/v SDS, ∼0.4% Bromophenol Blue (G250), and 40% v/v glycerol) and heated at 95 °C for 3 min. Samples were loaded into 0.75 mm 15% SDS-PAGE gels, and electrophoresis was carried out at a constant 120 V using 1X Tris-Glycine SDS running buffer (25 mM Tris base, 205 mM glycine, 3.45 mM SDS). Proteins were transferred to PVDF membranes using a wet transfer system with 1× Tris-Glycine transfer buffer (25 mM Tris base, 205 mM glycine, and 20% v/v methanol) at 85 V for 45 min. Membranes were blocked with 5% w/v BSA in 0.005% TBST (1.5M NaCl, 0.1M Tris–HCl pH 7.5, 0.005% v/v Triton X-100) at room temperature for 1 h. Primary antibodies were diluted in 5% BSA in TBST at the following ratios: mouse anti-UBA52 (rpL40, BioRad VMA00464), 1:2500; mouse anti-rpS3 (Proteintech 66046-1-Ig), 1:10000; rabbit anti-rpL3 (Proteintech 11005-1-AP), 1:20000; mouse anti-HSP90 (BD Transduction 610418), 1:5000; rabbit anti-rpL36 (Proteintech 15145-1-AP), 1:5000; rabbit anti-LC3B (Proteintech 18725-1-AP), 1:5000. Membranes were incubated with primary antibodies overnight at 4 °C, washed three times with 0.005% TBST, then incubated with HRP-linked anti-rabbit or anti-mouse secondary antibodies (1:5000 in 5% milk in 0.005% TBST) at room temperature for 1 h. Membranes were washed three times with 0.005% TBST, and imaged using ECL western blotting substrate and a Biorad Chemidoc imaging system.

### Polysome profiling and fractionation

Polysome profiling was performed as previously described^17^. Briefly, cells were treated with 100 µg mL^-1^ cycloheximide (MP Biomedicals) for 5 min before harvesting in cold PBS containing 100 µg mL^-1^ cycloheximide. Cells were pelleted by centrifugation at 300 × g for 3 min at 4 °C. Cell pellets were lysed in 200 µL ice-cold polysome lysis buffer (20 mM Tris–HCl pH 7.5, 200 mM NaCl, 10 mM MgCl_2_, 0.1% v/v Triton X-100, 1 mM DTT, 1X protease inhibitor cocktail, 100 µg mL^-1^ cycloheximide) by incubating on ice for 6 min and triturating through a 21-gauge needle five times. Nuclei and cell debris were removed by centrifugation at 10,000 × g for 5 min at 4 °C, and equal A_260_ units of lysates were loaded on a 10–50% w/v sucrose gradient made in polysome buffer (20 mM Tris–HCl pH 7.5, 150 mM NaCl, 10 mM MgCl_2_, 1 mM DTT) prepared using a Brandel gradient maker. Gradients were centrifuged for 2 h at 4 °C in a TH-641 rotor. After centrifugation, gradients were fractionated using a Brandel gradient fractionator with A_254_ profile monitoring by a Dataq data acquisition system. To purify 80S or polysome fractions for immunoblotting, native gel shift assays, or mass spectrometry, appropriate fractions were diluted with polysome buffer without sucrose at a 1:1 ratio and centrifuged at 30,000 rpm for 16 h at 4 °C in a 70.1 Ti rotor. Ribosome pellets were resuspended in 30 µL storage buffer (25 mM HEPES–KOH pH 7.5, 100 mM KOAc, 10 mM MgOAc_2_, 6% w/v sucrose, 1 mM DTT, and protease inhibitor cocktail) by gentle pipetting ∼250 times. Ribosome concentration was determined by A_260_ reading (1 A_260_ = 20nM ribosomes). For Polysome-Seq, RNA from heavy polysomes (fractions 8–11) was purified by phenol-chloroform extraction. 200 µL of each fraction was diluted 1:1 with RNase-free water, and RNA was extracted by phenol-chloroform and precipitated with ethanol. cDNA synthesis and quantitative PCR was performed as described above.

### Ribosome–RNA native gel shift

^32^P-labeled VSV RNAs were incubated with 80S ribosomes, purified from VSV-infected HEK293T cells, at a final concentration series of 25 nM, 50 nM, 75 nM, 100 nM, and 200 nM, in EMSA incubation buffer (25 mM HEPES–KOH pH 7.5, 100 mM KOAc, 2.5 mM MgOAc_2_, 0.42 mM spermidine, 2 mM DTT) at 37 °C for 10 min. Reactions were immediately subjected to electrophoresis on a 0.7% w/v native agarose gel 1× TBE buffer (89 mM Tris base, 89 mM boric acid, 2 mM EDTA pH 8.0) with 75 mM KCl at 100 V for 42 min on ice. The gel was fixed in a buffer containing 40% v/v acetic acid and 10% ethanol v/v, then dried at 75.5 °C for 2 hours. Following drying, the gel was exposed to a phosphor screen and imaged using an Amersham Typhoon IP system (Cytiva). For the competition assay, ^32^P-labeled VSV mRNA was mixed with 100 nM 80S ribosomes purified from VSV-infected HEK293T cells, with or without a 5-fold molar excess of unlabeled β-actin or VSV mRNA, using the same conditions as described above. For antibody inhibition, 100 nM purified 80S ribosomes was pre-incubated with 0.5 µg of UBA52 (rpL40)-targeting antibody or mouse IgG control (Invitrogen 02-6502) at room temperature for 10 min, followed by incubation with ^32^P-labeled VSV mRNAs as described above.

### Tandem Mass Tag-mass spectrometry (TMT-MS)

Mass spectrometry sample preparation. After collecting the appropriate fraction from a polysome profiling experiment, buffer exchange was performed using a modified SP3 protocol^78^. Briefly, ∼250 µg of Cytiva Speed-Bead Magnetic Carboxylate Modified Particles (65152105050250 and 4515210505250), mixed at a 1:1 ratio, were added to each sample. 100% ethanol was added to each sample to achieve a final ethanol concentration of at least 50%. Samples were incubated with gentle shaking for 15 min. Samples were washed three times with 80% ethanol. Protein was eluted from SP3 beads using 200 mM EPPS pH 8.5 containing Lys-C (Wako, 129-02541). Samples were digested overnight at room temperature with vigorous shaking. The next morning trypsin was added to each sample and further incubated for 6 hours at 37 ºC. Acetonitrile was added to each sample to achieve a final concentration of ∼33%. Each sample was labelled, in the presence of SP3 beads, with ∼62.5 µg of TMT reagents (ThermoFisher Scientific). Following confirmation of satisfactory labelling (>97%), excess TMT was quenched by addition of hydroxylamine to a final concentration of 0.3%. The full volume from each sample was pooled and acetonitrile was removed by vacuum centrifugation. The samples were acidified with formic acid and desalted by StageTip eluted into autosampler inserts (Thermo Scientific), dried in a speedvac and reconstituted with 5% Acetonitrile, 5% formic acid for LC-MS/MS analysis.

Liquid chromatography and tandem mass spectrometry. Two pairs of biologically independent samples of mock and VSV-infected 80S ribosomes were collected and analyzed together in a single mass spectrometry run. Data were collected on an Orbitrap Eclipse mass spectrometer coupled to a Proxeon NanoLC-1200 UHPLC (Thermo Fisher Scientific). The 100 µm capillary column was packed in-house with 35 cm of Accucore 150 resin (2.6 µm, 150Å; ThermoFisher Scientific). Data were acquired for 180 min per run. A FAIMS device was enabled during data collection and compensation voltages were set at −40V, −60V, and −80V^79^. MS1 scans were collected in the Orbitrap (resolution – 60,000; scan range – 400-1600 Th; automatic gain control (AGC) – 4× 10^5^; maximum ion injection time – automatic). MS2 scans were collected in the Orbitrap following higher-energy collision dissociation (HCD; resolution – 50,000; AGC – 250%; normalized collision energy – 36; isolation window – 0.5 Th; maximum ion injection time – 86 ms.

Mass Spectrometry Data Analysis. Database searching included all entries from the human UniProt Database (downloaded in May 2021). The database was concatenated with one composed of all protein sequences for that database in the reversed order^80^. Raw files were converted to mzXML, and monoisotopic peaks were re-assigned using Monocle^81^. Searches were performed with Comet^82^ using a 50-ppm precursor ion tolerance and fragment bin tolerance of 0.02. TMT labels on lysine residues and peptide N-termini (+229.1629 Da), as well as carbamidomethylation of cysteine residues (+57.021 Da) were set as static modifications, while oxidation of methionine residues (+15.995 Da) was set as a variable modification. Peptide-spectrum matches (PSMs) were adjusted to a 1% false discovery rate (FDR) using a linear discriminant after which proteins were assembled further to a final protein-level FDR of 1% analysis^83^. TMT reporter ion intensities were measured using a 0.003 Da window around the theoretical m/z for each reporter ion. Proteins were quantified by summing reporter ion counts across all matching PSMs. More specifically, reporter ion intensities were adjusted to correct for the isotopic impurities of the different TMT reagents according to manufacturer specifications. Peptides were filtered to exclude those with a summed signal-to-noise (SN) < 100 across all TMT channels and < 0.5 precursor isolation specificity. The signal-to-noise (S/N) measurements of peptides assigned to each protein were summed (for a given protein). To create the heat map analysis, only ribosomal proteins were included, and differential protein abundance was assessed using the limma package^84,85^ on log_2_-transformed values, with statistical significance was defined as an adjusted P-value < 0.05.

### Crosslinking mass spectrometry (XL-MS)

Samples were cross-linked as previously described^24^. Briefly, the cross-linking reaction was performed with 2 mM BS3 (Thermo) in 50 mM HEPES, 100 mM NaCl, pH 7.8 for 1 h at room temperature. Reactions were quenched with hydroxylamine to a final concentration of 50 mM. All samples were reduced for 1 h in 2% sodium deoxycholate and 10 mM tris(2-carboxyethyl)phosphine, alkylated with 20 mM iodoacetamide in the dark for 1h and quenched with 20 mM β-mercaptoethanol for 15 min. Samples were then processed using the SP3 method^86^ and digested with LysC(Wako) for 3 h and then trypsin (Promega) for 6 h, both at 1:30 enzyme:substrate ratio and 37 °C. Digested peptides were acidified with 10% TFA to pH ∼2, desalted using stage tips with Empore C18 SPE Extraction Disks (3M) and dried under vacuum.

Samples were reconstituted in 5% FA/5% acetonitrile and analyzed in the Orbitrap Eclipse Mass Spectrometer (Thermo Fisher Scientific) coupled to an EASY-nLC 1200 (Thermo Fisher Scientific) ultra-high pressure liquid chromatography pump, as well as a high-field asymmetric waveform ion mobility spectrometry (FAIMS) interface. Peptides were separated on an inhouse pulled 100-µm inner diameter column packed with 35 cm of Accucore C18 resin (2.6 µm, 150 Å, Thermo Fisher Scientific), using a gradient consisting of 5–35% (acetonitrile, 0.125% FA) over 180 min at ∼550 nL min^−1^. The instrument was operated in data-dependent mode. FTMS1 spectra were collected at a resolution of 60K, with an AGC target of 100% and a maximum injection time of 50 ms. The most intense ions were selected for tandem MS (MS/MS) for 1.5 s in top-speed mode, while switching among three FAIMS compensation voltages (CVs), −40, –60 and –80 V, in the same method. Precursors were filtered according to charge state (allowed 3 ≤ *z* ≤ 7) and monoisotopic peak assignment was turned on. Previously interrogated precursors were excluded using a dynamic exclusion window (120 s ± 10 ppm). MS2 precursors were isolated with a quadrupole mass filter set to a width of 0.7 *m*/*z* and analyzed by FTMS2, with the Orbitrap operating at 30K resolution, an nAGC target of 250% and a maximum injection time of 150 ms. Precursors were then fragmented by high-energy collision dissociation at 30% normalized collision energy.

Mass spectra were processed and searched using the PIXL search engine^24^. The sequence database contained 2549 proteins identified at 1% false discovery range (FDR) in a non-crosslinked search. For cross-linked search, precursor tolerance was set to 15 ppm and fragment ion tolerance to 10 ppm. Methionine oxidation and protein N-terminal acetylation were set as variable modifications in addition to mono-linked mass of +156.0786 for BS3. Cross-linker mass shift of +138.0681 was used for BS3. Top 100 most abundant proteins by total spectral counts were considered by PIXL for cross-linking. Matches were filtered to 1% FDR on the unique peptide level using linear discriminant features as previously described^24^.

### Structural modeling

Structural modeling of crosslinking sites was generated using ChimeraX-1.10^87^. The 80S ribosome model was based on PDB 4UG0 (core)^69^, with mRNA incorporated from PDB 8OZ0^88^ and P1 protein elements incorporated from PDB 8XSZ^89^. The three models were aligned and merged for visualization. The model of alternative rpL40 interaction with the ribosome was created using the Chai-1 webserver^68^ using rpS3, rpS10, rpL40, rpS30 and a partial sequence of 18S RNA, with the two rpL40-rpS3 cross-links as contact restraints with maximum distance of 20 Å. The resulting model was aligned with 4UG0 to position rpL40 on the full ribosome structure.

### Crosslinking-immunoblotting

80S ribosomes used for crosslinking-immunoblotting were isolated from HEK293T cells stably expressing His-rpS3 after mock or VSV-infection at MOI = 3 for 3 h. To perform crosslinking, 0.5 pmol of 80S ribosomes were incubated with DSP at a final concentration of 0.05 mM. The total reaction volume was adjusted to 10 µL with PBS, and the reaction was incubated at room temperature for 30 min. Crosslinking was then quenched by adding Tris–HCl pH 7.5 to a final concentration of 50 mM, followed by incubation at room temperature for 15 min. The crosslinked 80S ribosomes were dissociated by treatment with 4 M urea at 37 °C for 20 min. For His-rpS3 pulldown, the dissociated ribosomes were incubated with Ni-NTA beads in binding buffer (1 M urea, 50 mM Tris–HCl pH 7.5) at 4 °C for 1 h. After binding, the beads were washed three times with washing buffer (binding buffer supplemented with 30 mM imidazole). Beads were pelleted by centrifugation and denatured in SDS-loading buffer with or without β-mercaptoethanol as denoted. Samples were then subjected to SDS-PAGE and immunoblotting analysis.

### Microscopy and image analysis

For immunofluorescence assays, cells were fixed with 4% PFA in PBS for 15 min at room temperature and washed 3 times with PBS. Cells were then permeabilized with 0.2% TritonX-100 in PBS for 5 min at room temperature, washed 3 times with PBS, and blocked with 5% goat serum in PBS for 1 h. Primary antibody incubation against cleaved caspase 3 (1:100, Proteintech 25128-1-AP) was done overnight at 4 °C. Cells were then washed 3 times with PBS and incubated with Alexa Fluor-594 conjugated goat anti-Rabbit IgG (1:1000, Invitrogen A-11012) for 2 h. Slides were washed 3 times and mounted using VECTASHIELD Antifade Mounting Medium with DAPI. For microscopy of viruses expressing fluorescent markers, all images were acquired on an EVOS MA5000 at a 10× objective.

Fluorescence image processing and quantification was performed using FIJI/ImageJ software. For cell death assays using cleaved-caspase 3 (Alexa Fluor-594, RFP), background noise was uniformly subtracted in RFP images using the Subtract function in ImageJ (Process > Math) with a constant value of 55 across all images. Briefly, binary masks of each channel were created using ImageJ thresholding functions. The algorithm ‘intermodes’ was used for RFP channels and ‘moments’ for BFP channels. The same threshold was applied to all images within the respective color channel. For RFP images, binary masks were then overlayed onto the fluorescent image to determine positive signal. The integrated density for RFP and BFP (nuclei) channels was measured in FIJI/ImageJ using a region of interest (ROI) of 300 μm^2^. To account for differences in cell number within each ROI, RFP (cleaved-caspase 3) integrated density was normalized to BFP (nuclei) integrated density. Three separate biological replicates (36 ROIs each) were analyzed per experimental group.

For all viral GFP images, the Subtract Background function in FIJI/ImageJ (Process > Subtract Background; rolling ball radius = 100) was uniformly applied to remove uneven illumination and background noise. Additional brightness and contrast adjustments (Image > Adjust > Brightness/Contrast) to GFP images in Fig. 2E and Fig. S2C were done using identical parameters across all images within the same experiment. To normalize fluorescent intensity between images in Fig. 1J, the product of the gain, exposure time, and light source intensity of the off-scale image (MeV-GFP) was normalized to the product of the gain, exposure time, and light source intensity of the reference images (mock and VSV-GFP). The resulting scale factor of 2.09 was then applied to each pixel of the off-scale image via the Multiply function in FIJI/ImageJ (Process > Math > Multiply) to generate a normalized image for visualization only. All GFP adjustments were made for visualization purposes only.

### Bioinformatics

Libraries were prepared from total RNA or polysome-associated mRNAs using a Roche Kapa mRNA HyperPrep kit according to the manufacturer’s protocol on a Beckman Coulter Biomek i7 and sequenced on an Illumina NovaSeq X Plus at the Dana-Farber Cancer Institute Molecular Biology Core Facilities. For bioinformatics, sequencing reads were quality filtered and trimmed using Cutadapt^90^ and aligned to the human genome (GRCh38) using STAR^91^. The resulting alignments were converted to indexed BAM files with SAMtools^92^, and gene-level counts were generated using feature-Counts^93^. Differential gene enrichment between HeLa cells expressing WT or aN8 rpL40 under control or serum starvation was analyzed using DESeq2^94^, with P-value adjustments performed using fdrtool^95^, using one contrast to examine polysome-associated or transcription changes, and two contrast to examine translation efficiency changes. Transcripts regulated by ribosome remodeling were defined as having an adjusted P-value < 0.05 and a fold change > 2. Read distributions were visualized using wiggleplotR, and gene ontology was analyzed using DAVID^96^ with a significance threshold of P < 0.05.

**Figure S1.**
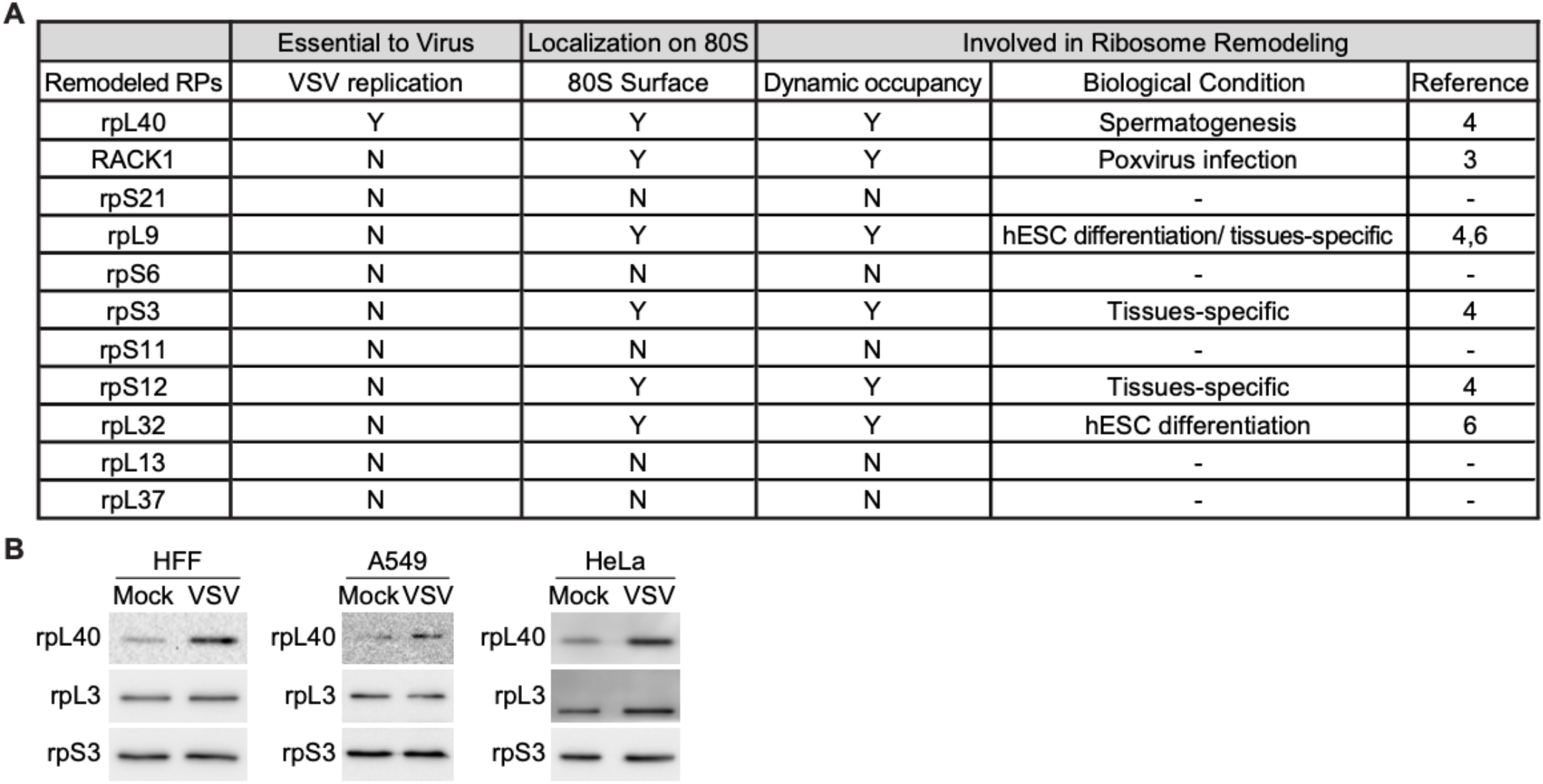
Increased rpL40 occupancy on ribosomes during VSV infection is conserved across cell types. **A**. Summary of ribosomal proteins significantly remodeled during VSV infection. **B**. Immunoblot of rpL40 levels on 80S ribosomes isolated from mock or VSV-infected primary human foreskin fibroblasts (HFF), human lung carcinoma A549, or human cervical cancer HeLa cells. The results are representative of *n* = 3 biologically independent samples.

**Figure S2.**
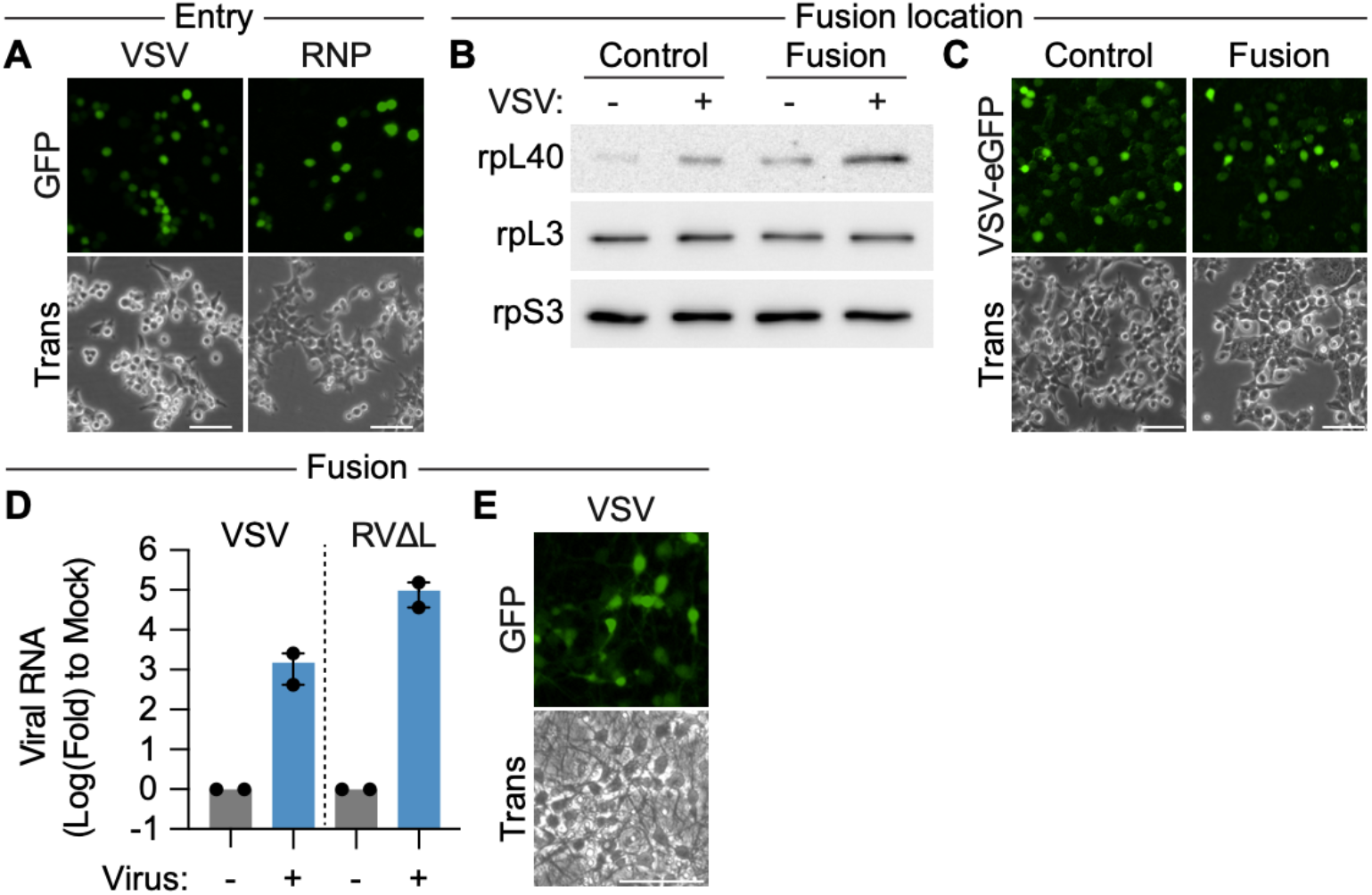
Measurement of viral replication upon perturbation of entry steps. **A**. Fluorescence microscopy of HEK293T cells infected with VSV-eGFP (MOI = 3, 4 hpi) or transfected with VSV-eGFP RNPs. Scale bar, 300 µm. Trans, transmitted light. **B**. Immunoblot of rpL40 levels on 80S ribosomes isolated from HEK293T where VSV entry occurs through a normal endosomal route (control) or via low pH-induced fusion at the plasma membrane. **C**. Fluorescence microscopy of HEK293T cells infected with VSV-eGFP through normal entry or fusion at the plasma membrane. Scale bar, 300 μm. **D**. Levels of viral genomic RNA levels upon infection of primary mouse cortical neurons with VSV or polymerase-deficient Rabies virus (RVΔL), as measured by qRT-PCR targeting of the *N* gene. Results are presented as mean fold change ± SEM relative to the corresponding mock control from *n* = 2 biologically independent samples. **E**. Fluorescence microscopy of primary mouse cortical neurons infected with VSV-eGFP (MOI 3, 4 h). Scale bar, 75 μm. Results of (E) are representative of *n* = 2 biologically independent samples. Results of (A–C) are representative of *n* = 3 biologically independent samples.

**Figure S3.**
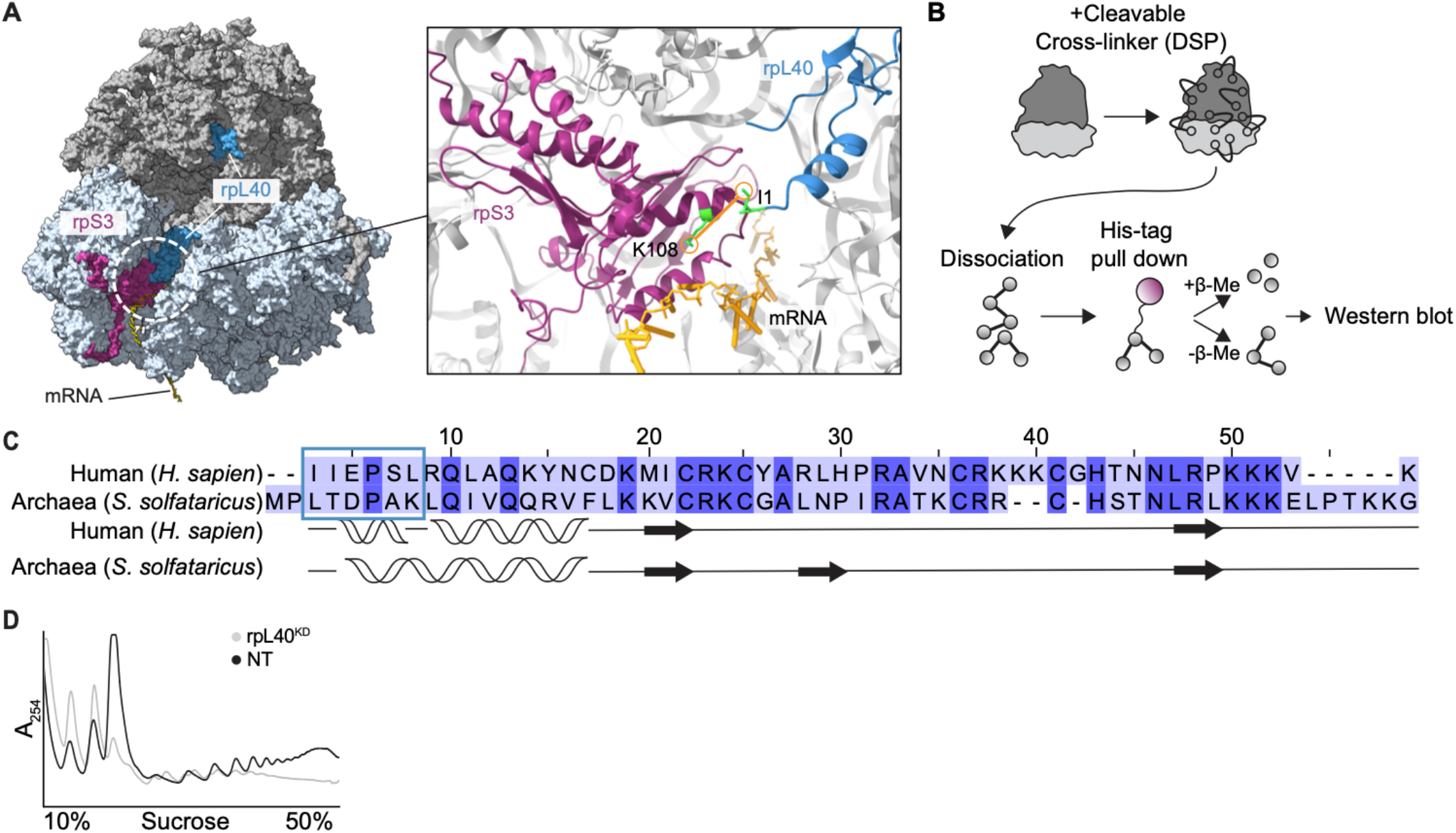
Structural modeling and sequence conservation analysis of rpL40. **A**. Chai-1^68^ structural modeling with crosslinking restraints of the predicted interaction site of rpL40 with the small ribosomal subunit, merged with the human 80S ribosome model (PDB 4UG0)^69^ for visualization. **B**. Schematic of crosslinking-immunoblot workflow. **C**. Sequence alignment and structural features of human and archaea rpL40. Sequence alignment is colored by % identity, with dark purple indicating identical residues. **D**. Polysome profiles from HeLa cells transfected with non-targeting (NT) or rpL40-targeting (rpL40^KD^) siRNA, with matching axes to **Fig. 3G**. Results of (D) are representative of *n* = 3 biologically independent samples.

**Figure S4.**
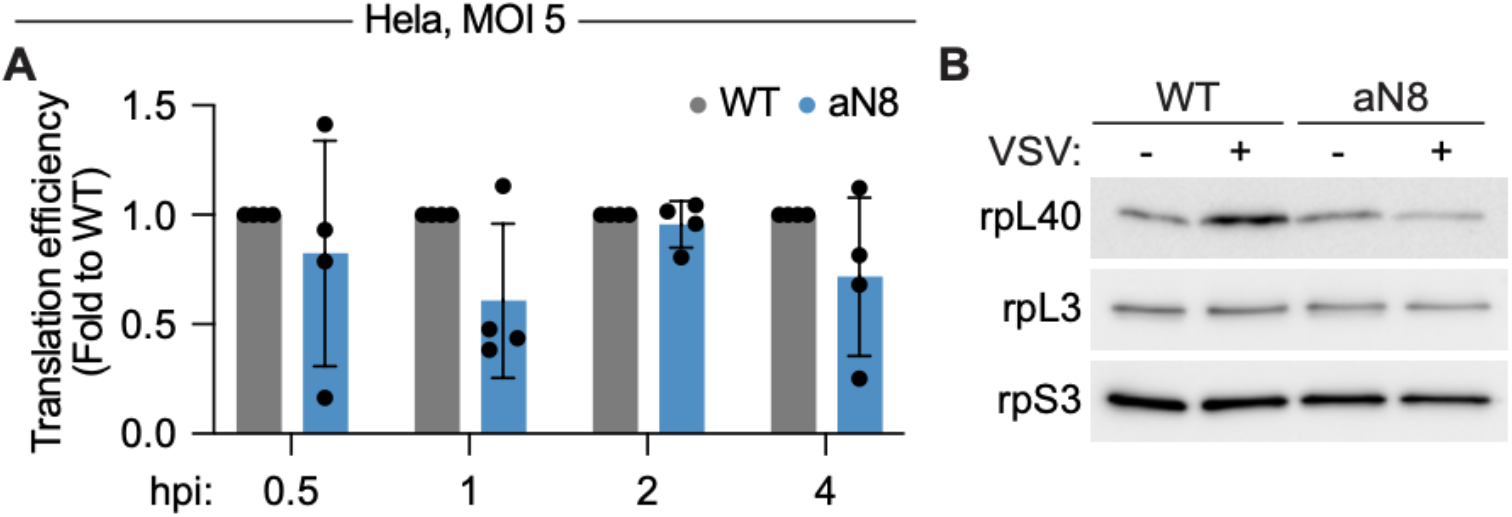
Ribosome remodeling is not critical for viral infection under permissive conditions. **A**. VSV-*Fluc* mRNA translation efficiency during a high MOI infection timecourse in HeLa cells exclusively expressing wild type (WT) or aN8 (occupancy mutant) rpL40 (MOI = 5). The results are presented as mean fold change ± SD in translation efficiency relative to WT cells, *n* = 3 biologically independent samples. **B**. Immunoblot of rpL40 levels on 80S ribosomes isolated from mock or VSV-infected A549 cells exclusively expressing WT or aN8 rpL40. The result is representative of *n* = 3 biologically independent samples.

**Figure S5.**
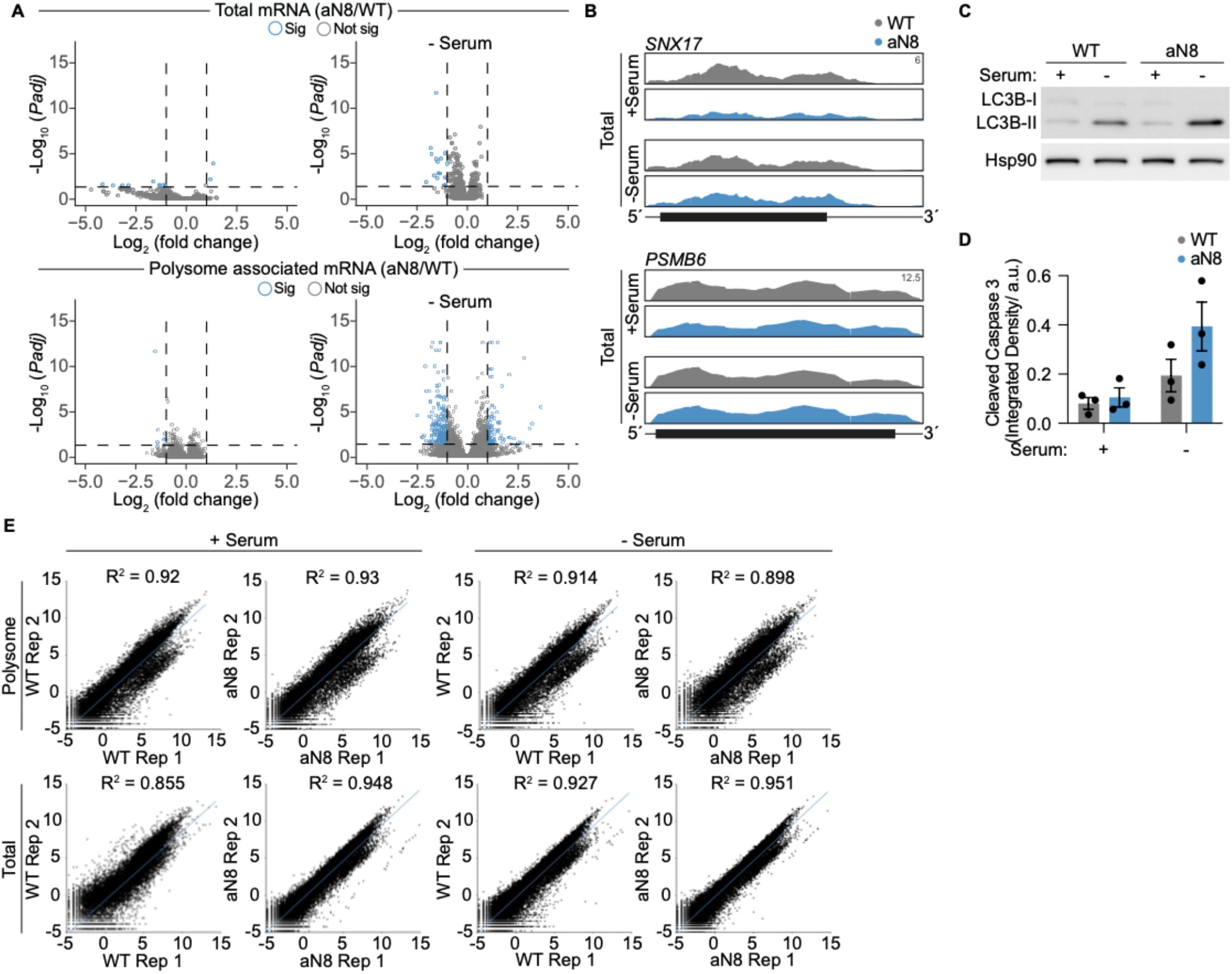
Ribosome remodeling is required for translation adaptation during serum starvation. **A**. Volcano plot comparing the fold change in cytoplasmic transcript (top) or polysome-associated (bottom) mRNA levels and adjusted *P* value in aN8 occupancy mutant versus wild type rpL40-expressing HeLa cells under control (left) or serum starvation for 6 h (right). **B**. Read mapping to *SNX17* and *PSMB6* from RNA-Seq under control or serum starvation conditions. The annotated y-axis maximum is equivalent for all samples. Transcript architecture is represented as follows: UTRs (thin line), coding regions (thick line). **C**. Immunoblot of LC3B cleavage in cytoplasmic extracts from WT versus aN8 rpL40 mutant HeLa cell lines under control or serum starvation conditions. **D**. Quantification of cleaved-caspase 3 levels shown in **Fig. 5I**. RFP intensity is plotted as mean ± SEM from *n* = 3 biologically independent samples. **E**. Reproducibility scatterplots of CPMs from total RNA-Seq and Polysome-Seq data. Results of (A) and (B) are from *n* = 2 biologically independent samples. Results of (C) are representative of *n* = 3 biologically independent samples.

## Supplementary: Table legends

**Table 1. Levels of ribosomal proteins in 80S ribosomes under mock or VSV-infected conditions, identified by TMT-MS**. A list of the summed signal-to-noise (S/N) ratios, fold change of 80S-association between mock and VSV-infected conditions, and statistical results for each ribosomal protein. Two independent biological replicates were combined in a single mass spectrometry acquisition.

**Table 2. Cross-linked sites identified by 80S ribosome XL-MS**. A list of rpL40 cross-linked peptides identified from 80S ribosomes purified from VSV-infected cells (Sheet 1). For each site, the identified peptides, the residue number in the full-length protein (global position of cross-linked protein), the position within the detected peptide sequence (local position of cross-linked peptide), and biological replicates in which they were identified. All identified crosslinked peptides from 80S ribosome XL-MS from 2 biological replicates are in Sheets 2 and 3.

**Table 3. Cellular mRNAs translationally regulated by increased rpL40 ribosomal occupancy**. Lists of gene with decreased translation efficiency in cells expressing aN8 versus WT rpL40, under complete media or serum starvation.

**Table S1. Table of oligonucleotides**. List of primers used for cloning and qRT-PCR.

## References

1 Klinge, S., Voigts-Hoffmann, F., Leibundgut, M. & Ban, N. Atomic structures of the eukaryotic ribosome. Trends Biochem Sci 37, 189–198 (2012). 10.1016/j.tibs.2012.02.007

2 Dopler, A. et al. P-stalk ribosomes act as master regulators of cytokine-mediated processes. Cell 187, 6981–6993 e6923 (2024). 10.1016/j.cell.2024.09.039

3 Khalatyan, N. et al. Ribosome customization and functional diversification among P-stalk proteins regulate late poxvirus protein synthesis. Cell Rep 44, 115119 (2025). 10.1016/j.celrep.2024.115119

4 Li, H. et al. A male germ-cell-specific ribosome controls male fertility. Nature 612, 725–731 (2022). 10.1038/s41586-022-05508-0

5 Shi, Z. et al. Heterogeneous Ribosomes Preferentially Translate Distinct Subpools of mRNAs Genome-wide. Mol Cell 67, 71–83 e77 (2017). 10.1016/j.molcel.2017.05.021

6 Genuth, N. R. et al. A stem cell roadmap of ribosome heterogeneity reveals a function for RPL10A in mesoderm production. Nat Commun 13, 5491 (2022). 10.1038/s41467-022-33263-3

7 Yang, Y. M. & Karbstein, K. The chaperone Tsr2 regulates Rps26 release and reincorporation from mature ribosomes to enable a reversible, ribosome-mediated response to stress. Sci Adv 8, eabl4386 (2022). 10.1126/sciadv.abl4386

8 Landry, D. M., Hertz, M. I. & Thompson, S. R. RPS25 is essential for translation initiation by the Dicistroviridae and hepatitis C viral IRESs. Genes Dev 23, 2753–2764 (2009). 10.1101/gad.1832209

9 Majzoub, K. et al. RACK1 controls IRES-mediated translation of viruses. Cell 159, 1086–1095 (2014). 10.1016/j.cell.2014.10.041

10 Schubert, K. et al. Author Correction: SARS-CoV-2 Nsp1 binds the ribosomal mRNA channel to inhibit translation. Nat Struct Mol Biol 27, 1094 (2020). 10.1038/s41594-020-00533-x

11 Tidu, A. et al. The viral protein NSP1 acts as a ribosome gatekeeper for shutting down host translation and fostering SARS-CoV-2 translation. RNA 27, 253–264 (2020). 10.1261/rna.078121.120

12 Havkin-Solomon, T. et al. Selective translational control of cellular and viral mRNAs by RPS3 mRNA binding. Nucleic Acids Res 51, 4208–4222 (2023). 10.1093/nar/gkad269

13 Temin, H. M. & Mizutani, S. RNA-dependent DNA polymerase in virions of Rous sarcoma virus. Nature 226, 1211–1213 (1970). 10.1038/2261211a0

14 Berget, S. M., Moore, C. & Sharp, P. A. Spliced segments at the 5’ terminus of adenovirus 2 late mRNA. Proceedings of the National Academy of Sciences of the United States of America 74, 3171–3175 (1977). 10.1073/pnas.74.8.3171

15 Sarnow, P. Translation of glucose-regulated protein 78/immunoglobulin heavy-chain binding protein mRNA is increased in poliovirus-infected cells at a time when cap-dependent translation of cellular mRNAs is inhibited. Proc Natl Acad Sci U S A 86, 5795–5799 (1989). 10.1073/pnas.86.15.5795

16 Finley, D., Bartel, B. & Varshavsky, A. The tails of ubiquitin precursors are ribosomal proteins whose fusion to ubiquitin facilitates ribosome biogenesis. Nature 338, 394–401 (1989). 10.1038/338394a0

17 Lee, A. S., Burdeinick-Kerr, R. & Whelan, S. P. A ribosome-specialized translation initiation pathway is required for cap-dependent translation of vesicular stomatitis virus mRNAs. Proc Natl Acad Sci U S A 110, 324–329 (2013). 10.1073/pnas.1216454109

18 Cureton, D. K., Massol, R. H., Saffarian, S., Kirchhausen, T. L. & Whelan, S. P. Vesicular stomatitis virus enters cells through vesicles incompletely coated with clathrin that depend upon actin for internalization. PLoS Pathog 5, e1000394 (2009). 10.1371/journal.ppat.1000394

19 Slough, M. M., Chandran, K. & Jangra, R. K. Two Point Mutations in Old World Hantavirus Glycoproteins Afford the Generation of Highly Infectious Recombinant Vesicular Stomatitis Virus Vectors. mBio 10 (2019). 10.1128/mBio.02372-18

20 Brown, K. S., Safronetz, D., Marzi, A., Ebihara, H. & Feldmann, H. Vesicular stomatitis virus-based vaccine protects hamsters against lethal challenge with Andes virus. J Virol 85, 12781–12791 (2011). 10.1128/JVI.00794-11

21 Jin, L. et al. Third-generation rabies viral vectors allow nontoxic retrograde targeting of projection neurons with greatly increased efficiency. Cell Rep Methods 3, 100644 (2023). 10.1016/j.crmeth.2023.100644

22 Klinge, S., Voigts-Hoffmann, F., Leibundgut, M., Arpagaus, S. & Ban, N. Crystal structure of the eukaryotic 60S ribosomal subunit in complex with initiation factor 6. Science 334, 941–948 (2011). 10.1126/science.1211204

23 Yu, C. & Huang, L. Cross-Linking Mass Spectrometry: An Emerging Technology for Interactomics and Structural Biology. Anal Chem 90, 144–165 (2018). 10.1021/acs.analchem.7b04431

24 Mintseris, J. & Gygi, S. P. High-density chemical cross-linking for modeling protein interactions. Proc Natl Acad Sci U S A 117, 93–102 (2020). 10.1073/pnas.1902931116

25 Dong, J. et al. Rps3/uS3 promotes mRNA binding at the 40S ribosome entry channel and stabilizes preinitiation complexes at start codons. Proc Natl Acad Sci U S A 114, E2126–E2135 (2017). 10.1073/pnas.1620569114

26 Babaylova, E., Malygin, A., Gopanenko, A., Graifer, D. & Karpova, G. Tetrapeptide 60-63 of human ribosomal protein uS3 is crucial for translation initiation. Biochim Biophys Acta Gene Regul Mech 1862, 194411 (2019). 10.1016/j.bbagrm.2019.194411

27 Wang, J., Zhou, J., Yang, Q. & Grayhack, E. J. Multi-protein bridging factor 1(Mbf1), Rps3 and Asc1 prevent stalled ribosomes from frameshifting. Elife 7 (2018). 10.7554/eLife.39637

28 Wu, B. et al. Solution structure of ribosomal protein L40E, a unique C4 zinc finger protein encoded by archaeon Sulfolobus solfataricus. Protein Sci 17, 589–596 (2008). 10.1110/ps.073273008

29 Barr, J. N. & Wertz, G. W. Polymerase slippage at vesicular stomatitis virus gene junctions to generate poly(A) is regulated by the upstream 3’-AUAC-5’ tetranucleotide: implications for the mechanism of transcription termination. J Virol 75, 6901–6913 (2001). 10.1128/JVI.75.15.6901-6913.2001

30 Abraham, G., Rhodes, D. P. & Banerjee, A. K. The 5’ terminal structure of the methylated mRNA synthesized in vitro by vesicular stomatitis virus. Cell 5, 51–58 (1975). 10.1016/0092-8674(75)90091-4

31 Walsh, D. & Mohr, I. Viral subversion of the host protein synthesis machinery. Nat Rev Microbiol 9, 860–875 (2011). 10.1038/nrmicro2655

32 Kozak, M. A short leader sequence impairs the fidelity of initiation by eukaryotic ribosomes. Gene Expr 1, 111–115 (1991).

33 Baltimore, D., Huang, A. S. & Stampfer, M. Ribonucleic acid synthesis of vesicular stomatitis virus, II. An RNA polymerase in the virion. Proc Natl Acad Sci U S A 66, 572–576 (1970). 10.1073/pnas.66.2.572

34 Bushell, M. & Sarnow, P. Hijacking the translation apparatus by RNA viruses. J Cell Biol 158, 395–399 (2002). 10.1083/jcb.200205044

35 Franz, K. M. & Kagan, J. C. Innate Immune Receptors as Competitive Determinants of Cell Fate. Mol Cell 66, 750–760 (2017). 10.1016/j.molcel.2017.05.009

36 Blach-Olszewska, Z., Halasa, J., Matej, H. & Cembrzynska-Nowak, M. Why HeLa cells do not produce interferon? Arch Immunol Ther Exp (Warsz) 25, 683–691 (1977).

37 Landau, L. M. et al. pLxIS-containing domains are biochemically flexible regulators of interferons and metabolism. Mol Cell 84, 2436–2454 e2410 (2024). 10.1016/j.molcel.2024.05.030

38 Neidermyer, W. J., Jr. & Whelan, S. P. J. Global analysis of polysome-associated mRNA in vesicular stomatitis virus infected cells. PLoS Pathog 15, e1007875 (2019). 10.1371/journal.ppat.1007875

39 Krishnamurthy, S., Takimoto, T., Scroggs, R. A. & Portner, A. Differentially regulated interferon response determines the outcome of Newcastle disease virus infection in normal and tumor cell lines. J Virol 80, 5145–5155 (2006). 10.1128/JVI.02618-05

40 Tisoncik, J. R. et al. Into the eye of the cytokine storm. Microbiol Mol Biol Rev 76, 16–32 (2012). 10.1128/MMBR.05015-11

41 Foo, J., Bellot, G., Pervaiz, S. & Alonso, S. Mitochondria-mediated oxidative stress during viral infection. Trends Microbiol 30, 679–692 (2022). 10.1016/j.tim.2021.12.011

42 Le Sage, V., Cinti, A., Amorim, R. & Mouland, A. J. Adapting the Stress Response: Viral Subversion of the mTOR Signaling Pathway. Viruses 8 (2016). 10.3390/v8060152

43 Pakos-Zebrucka, K. et al. The integrated stress response. EMBO Rep 17, 1374–1395 (2016). 10.15252/embr.201642195

44 Weitzman, M. D. & Fradet-Turcotte, A. Virus DNA Replication and the Host DNA Damage Response. Annu Rev Virol 5, 141–164 (2018). 10.1146/annurev-virology-092917-043534

45 Wang, X. & Chen, X. J. A cytosolic network suppressing mitochondria-mediated proteostatic stress and cell death. Nature 524, 481–484 (2015). 10.1038/nature14859

46 Park, J. et al. SMYD5 methylation of rpL40 links ribosomal output to gastric cancer. Nature 632, 656–663 (2024). 10.1038/s41586-024-07718-0

47 Tripathi, M. et al. ESRRA (estrogen related receptor, alpha) induces ribosomal protein RPLP1-mediated adaptive hepatic translation during prolonged starvation. Autophagy 21, 1283–1297 (2025). 10.1080/15548627.2025.2465183

48 Sardiello, M. et al. A gene network regulating lysosomal biogenesis and function. Science 325, 473–477 (2009). 10.1126/science.1174447

49 Zhou, C. et al. Recycling of autophagosomal components from autolysosomes by the recycler complex. Nat Cell Biol 24, 497–512 (2022). 10.1038/s41556-022-00861-8

50 Tamai, K. et al. Role of Hrs in maturation of autophagosomes in mammalian cells. Biochem Biophys Res Commun 360, 721–727 (2007). 10.1016/j.bbrc.2007.06.105

51 Filimonenko, M. et al. Functional multivesicular bodies are required for autophagic clearance of protein aggregates associated with neuro-degenerative disease. J Cell Biol 179, 485–500 (2007). 10.1083/jcb.200702115

52 Ishimaru, K. et al. Sphingosine kinase-2 prevents macrophage cholesterol accumulation and atherosclerosis by stimulating autophagic lipid degradation. Sci Rep 9, 18329 (2019). 10.1038/s41598-019-54877-6

53 He, C. & Klionsky, D. J. Regulation mechanisms and signaling pathways of autophagy. Annu Rev Genet 43, 67–93 (2009). 10.1146/annurev-genet-102808-114910

54 Genuth, N. R. & Barna, M. The Discovery of Ribosome Heterogeneity and Its Implications for Gene Regulation and Organismal Life. Mol Cell 71, 364–374 (2018). 10.1016/j.molcel.2018.07.018

55 Ferretti, M. B. & Karbstein, K. Does functional specialization of ribosomes really exist? RNA 25, 521–538 (2019). 10.1261/rna.069823.118

56 Mauro, V. P. & Matsuda, D. Translation regulation by ribosomes: Increased complexity and expanded scope. RNA Biol 13, 748–755 (2016). 10.1080/15476286.2015.1107701

57 Fernandez-Pevida, A., Rodriguez-Galan, O., Diaz-Quintana, A., Kressler, D. & de la Cruz, J. Yeast ribosomal protein L40 assembles late into precursor 60 S ribosomes and is required for their cytoplasmic maturation. J Biol Chem 287, 38390–38407 (2012). 10.1074/jbc.M112.400564

58 Ohmayer, U. et al. Studies on the assembly characteristics of large subunit ribosomal proteins in S. cerevisae. PLoS One 8, e68412 (2013). 10.1371/journal.pone.0068412

59 Graifer, D., Malygin, A., Zharkov, D. O. & Karpova, G. Eukaryotic ribosomal protein S3: A constituent of translational machinery and an extraribosomal player in various cellular processes. Biochimie 99, 8–18 (2014). 10.1016/j.biochi.2013.11.001

60 Schubert, K. et al. SARS-CoV-2 Nsp1 binds the ribosomal mRNA channel to inhibit translation. Nat Struct Mol Biol 27, 959–966 (2020). 10.1038/s41594-020-0511-8

61 Yuan, S. et al. Nonstructural Protein 1 of SARS-CoV-2 Is a Potent Pathogenicity Factor Redirecting Host Protein Synthesis Machinery toward Viral RNA. Mol Cell 80, 1055–1066 e1056 (2020). 10.1016/j.molcel.2020.10.034

62 Jha, S. et al. Trans-kingdom mimicry underlies ribosome customization by a poxvirus kinase. Nature 546, 651–655 (2017). 10.1038/nature22814

63 Pelletier, J. & Sonenberg, N. Internal initiation of translation of eukaryotic mRNA directed by a sequence derived from poliovirus RNA. Nature 334, 320–325 (1988). 10.1038/334320a0

64 Wu, L. G. & Chan, C. Y. Membrane transformations of fusion and budding. Nat Commun 15, 21 (2024). 10.1038/s41467-023-44539-7

65 Kast, D. J. & Dominguez, R. The Cytoskeleton-Autophagy Connection. Curr Biol 27, R318–R326 (2017). 10.1016/j.cub.2017.02.061

66 Kast, D. J., Zajac, A. L., Holzbaur, E. L., Ostap, E. M. & Dominguez, R. WHAMM Directs the Arp2/3 Complex to the ER for Autophagosome Biogenesis through an Actin Comet Tail Mechanism. Curr Biol 25, 1791–1797 (2015). 10.1016/j.cub.2015.05.042

67 Schultz, N., Hamra, F. K. & Garbers, D. L. A multitude of genes expressed solely in meiotic or postmeiotic spermatogenic cells offers a myriad of contraceptive targets. Proc Natl Acad Sci U S A 100, 12201–12206 (2003). 10.1073/pnas.1635054100

68 Discovery, C. et al. Chai-1: Decoding the molecular interactions of life. bioRxiv, 2024.2010.2010.615955 (2024). 10.1101/2024.10.10.615955

69 Khatter, H., Myasnikov, A. G., Natchiar, S. K. & Klaholz, B. P. Structure of the human 80S ribosome. Nature 520, 640–645 (2015). 10.1038/nature14427

70 Gu, X. et al. The midnolin-proteasome pathway catches proteins for ubiquitination-independent degradation. Science 381, eadh5021 (2023). 10.1126/science.adh5021

71 Duprex, W. P., McQuaid, S., Hangartner, L., Billeter, M. A. & Rima, B. K. Observation of measles virus cell-to-cell spread in astrocytoma cells by using a green fluorescent protein-expressing recombinant virus. J Virol 73, 9568–9575 (1999). 10.1128/JVI.73.11.9568-9575.1999

72 Whelan, S. P., Barr, J. N. & Wertz, G. W. Identification of a minimal size requirement for termination of vesicular stomatitis virus mRNA: implications for the mechanism of transcription. J Virol 74, 8268–8276 (2000). 10.1128/jvi.74.18.8268-8276.2000

73 Whelan, S. P. & Wertz, G. W. Regulation of RNA synthesis by the genomic termini of vesicular stomatitis virus: identification of distinct sequences essential for transcription but not replication. J Virol 73, 297–306 (1999). 10.1128/JVI.73.1.297-306.1999

74 Lamper, A. M., Fleming, R. H., Ladd, K. M. & Lee, A. S. Y. A phosphorylation-regulated eIF3d translation switch mediates cellular adaptation to metabolic stress. Science 370, 853–856 (2020). 10.1126/science.abb0993

75 Kleinfelter, L. M. et al. Haploid Genetic Screen Reveals a Profound and Direct Dependence on Cholesterol for Hantavirus Membrane Fusion. mBio 6, e00801 (2015). 10.1128/mBio.00801-15

76 Whelan, S. P. & Wertz, G. W. Transcription and replication initiate at separate sites on the vesicular stomatitis virus genome. Proc Natl Acad Sci U S A 99, 9178–9183 (2002). 10.1073/pnas.152155599

77 Lee, A. S., Kranzusch, P. J. & Cate, J. H. eIF3 targets cell-proliferation messenger RNAs for translational activation or repression. Nature 522, 111–114 (2015). 10.1038/nature14267

78 Hughes, C. S. et al. Single-pot, solid-phase-enhanced sample preparation for proteomics experiments. Nat Protoc 14, 68–85 (2019). 10.1038/s41596-018-0082-x

79 Schweppe, D. K. et al. Characterization and Optimization of Multiplexed Quantitative Analyses Using High-Field Asymmetric-Wave-form Ion Mobility Mass Spectrometry. Anal Chem 91, 4010–4016 (2019). 10.1021/acs.analchem.8b05399

80 Elias, J. E. & Gygi, S. P. Target-decoy search strategy for increased confidence in large-scale protein identifications by mass spectrometry. Nat Methods 4, 207–214 (2007). 10.1038/nmeth1019

81 Rad, R. et al. Improved Monoisotopic Mass Estimation for Deeper Proteome Coverage. J Proteome Res 20, 591–598 (2021). 10.1021/acs.jproteome.0c00563

82 Eng, J. K., Jahan, T. A. & Hoopmann, M. R. Comet: an open-source MS/MS sequence database search tool. Proteomics 13, 22–24 (2013). 10.1002/pmic.201200439

83 Huttlin, E. L. et al. A tissue-specific atlas of mouse protein phosphorylation and expression. Cell 143, 1174–1189 (2010). 10.1016/j.cell.2010.12.001

84 Smyth, G. K. Linear models and empirical bayes methods for assessing differential expression in microarray experiments. Stat Appl Genet Mol Biol 3, Article3 (2004). 10.2202/1544-6115.1027

85 Ritchie, M. E. et al. limma powers differential expression analyses for RNA-sequencing and microarray studies. Nucleic Acids Res 43, e47 (2015). 10.1093/nar/gkv007

86 Moggridge, S., Sorensen, P. H., Morin, G. B. & Hughes, C. S. Extending the Compatibility of the SP3 Paramagnetic Bead Processing Approach for Proteomics. J Proteome Res 17, 1730–1740 (2018). 10.1021/acs.jproteome.7b00913

87 Pettersen, E. F. et al. UCSF ChimeraX: Structure visualization for researchers, educators, and developers. Protein Sci 30, 70–82 (2021). 10.1002/pro.3943

88 Brito Querido, J. et al. The structure of a human translation initiation complex reveals two independent roles for the helicase eIF4A. Nat Struct Mol Biol 31, 455–464 (2024). 10.1038/s41594-023-01196-0

89 Li, X. et al. Structural basis for differential inhibition of eukaryotic ribosomes by tigecycline. Nat Commun 15, 5481 (2024). 10.1038/s41467-024-49797-7

90 Martin, M. Cutadapt removes adapter sequences from high-throughput sequencing reads. 2011 17, 3 (2011). 10.14806/ej.17.1.200

91 Dobin, A. et al. STAR: ultrafast universal RNA-seq aligner. Bioinformatics 29, 15–21 (2013). 10.1093/bioinformatics/bts635

92 Li, H. et al. The Sequence Alignment/Map format and SAMtools. Bio-informatics 25, 2078–2079 (2009). 10.1093/bioinformatics/btp352

93 Liao, Y., Smyth, G. K. & Shi, W. featureCounts: an efficient general purpose program for assigning sequence reads to genomic features. Bioinformatics 30, 923–930 (2014). 10.1093/bioinformatics/btt656

94 Love, M. I., Huber, W. & Anders, S. Moderated estimation of fold change and dispersion for RNA-seq data with DESeq2. Genome Biol 15, 550 (2014). 10.1186/s13059-014-0550-8

95 Strimmer, K. fdrtool: a versatile R package for estimating local and tail area-based false discovery rates. Bioinformatics 24, 1461–1462 (2008). 10.1093/bioinformatics/btn209

96 Dennis, G. et al. DAVID: Database for Annotation, Visualization, and Integrated Discovery. Genome Biology 4, R60 (2003). 10.1186/gb-2003-4-9-r60

